# Structural properties of individual instances predict human effort and performance on an NP-Hard problem

**DOI:** 10.1101/405449

**Authors:** Juan Pablo Franco, Nitin Yadav, Peter Bossaerts, Carsten Murawski

## Abstract

Life presents us with decisions of varying degrees of difficulty. Many of them are NP-hard, that is, they are computationally intractable. Two important questions arise: which properties of decisions drive extreme computational hardness and what are the effects of these properties on human-decision making? Here, we postulate that we can study the effects of computational complexity on human decision-making by studying the mathematical properties of individual instances of NP-hard problems. We draw on prior work in computational complexity theory, which suggests that computational difficulty can be characterized based on the features of instances of a problem. This study is the first to apply this approach to human decision-making. We measured hardness, first, based on typical-case complexity (TCC), a measure of average complexity of a random ensemble of instances, and, second, based on instance complexity (IC), a measure that captures the hardness of a single instance of a problem, regardless of the ensemble it came from. We tested the relation between these measures and (i) decision quality as well as (ii) time expended in a decision, using two variants of the 0-1 knapsack problem, a canonical and ubiquitous computational problem. We show that participants expended more time on instances with higher complexity but that decision quality was lower in those instances. These results suggest that computational complexity is an inherent property of the instances of a problem, which affect human and other kinds of computers.

---

Life requires us to make complex decisions with limited cognitive resources. Many of the problems humans face are known to be computationally intractable in the sense that the time taken even by the most powerful digital computers to find a solution quickly grows to levels that makes solving these problems infeasible. Everyday life examples of deceptively simple yet actually hard problems include attention gating, task scheduling, shopping, routing, bin packing, and game play (1, 2). Behind the surface of these routine jobs lurk NP-hard problems such as the traveling salesperson problem, the knapsack problem, the Hamiltonian circuit problem, the graph coloring problem, and the K-SAT problems (boolean satisfiability problems). These problems are considered intractable because there are no known algorithms that can always find the solution with a number of computations that does not increase faster than a polynomial in the size of the instance at hand (3).

An important open question is in what ways computational complexity affects human cognition and decision-making. It has been shown that some task-dependent and some algorithm-dependent metrics of computational complexity predict human effort and performance in cognitive tasks including decisions (4–10). This approach is problematic, though, because these metrics are not readily generalizable to other problems and we cannot usually observe which algorithms people use when solving a cognitive problem.

Recently, it has been shown that computational difficulty is related to generic mathematical properties of instances of computational problems, independent of the algorithm employed (3, 11, 12). It is not known at this point to what extent these properties affect human behavior. If they do, it would strongly suggest that computational complexity is an inherent property of instances of computational problems (13, 14). This is the question we address here. More specifically, we explore whether generic problem features predict human effort and performance in a canonical decision task. We investigated how the mathematical properties of individual instances of the the knapsack problem affect human decision quality and time-on-task. We studied both the *decision variant* as well as the *optimization variant* of the 0-1 knapsack problem. In both variants, a set of items *I* is presented; each item has a weight *w* and a value *v*. In the *decision variant*, the task is to decide whether there exists a subset *A* of items from the set *I* for which (1) the sum of weights (∑_*i*∈*A*_*w*_*i*_) is lower or equal to a given capacity *c* and (2) the sum of values (∑_*i*∈*A*_ *v*_*i*_) is at least as high as a given target profit *p*. In the related *optimization variant*, the aim is to select the items that maximize the the sum of values (∑_*i∈A*_ *v*_*i*_) without exceeding the knapsack’s capacity (∑_*i*∈*A*_ *w*_*i*_ ≤*c*). Both variants are NP-hard (15). In our study, participants were asked to solve a number of random instances of both the decision variant and the optimization variant of the problem.

The knapsack problem is a canonical computational problem whose mathematical structure resembles that of many theories of decision-making such as utility maximization (16) and satisficing (17). However, its relevance extends beyond decision theory. The problem manifests itself in everyday life tasks involving choice of stimuli to attend to, budgeting and time management, portfolio optimization, intellectual discovery as well as in industrial applications such as the cargo business (15, 18). The problem has also been associated with symptoms of particular mental disorders, in particular, attention-deficit/hyperactivity disorder (19, 20).

We consider two metrics that generically capture instance hardness for digital computers tasked with finding the solution. First, we exploit findings on typical-case complexity (TCC), a popular approach for studying intrinsic hardness of random ensembles of instances of NP-hard problems (11, 12, 21). It has been shown that there exists a topology with which one can predict the average number of computations needed to find the solution of an instance (12, 22–27) (Fig 1a). We conjectured that TCC would predict performance and time-on-task for humans. We tested this in both the decision and optimization variants of the knapsack problem.

**Fig. 1.**
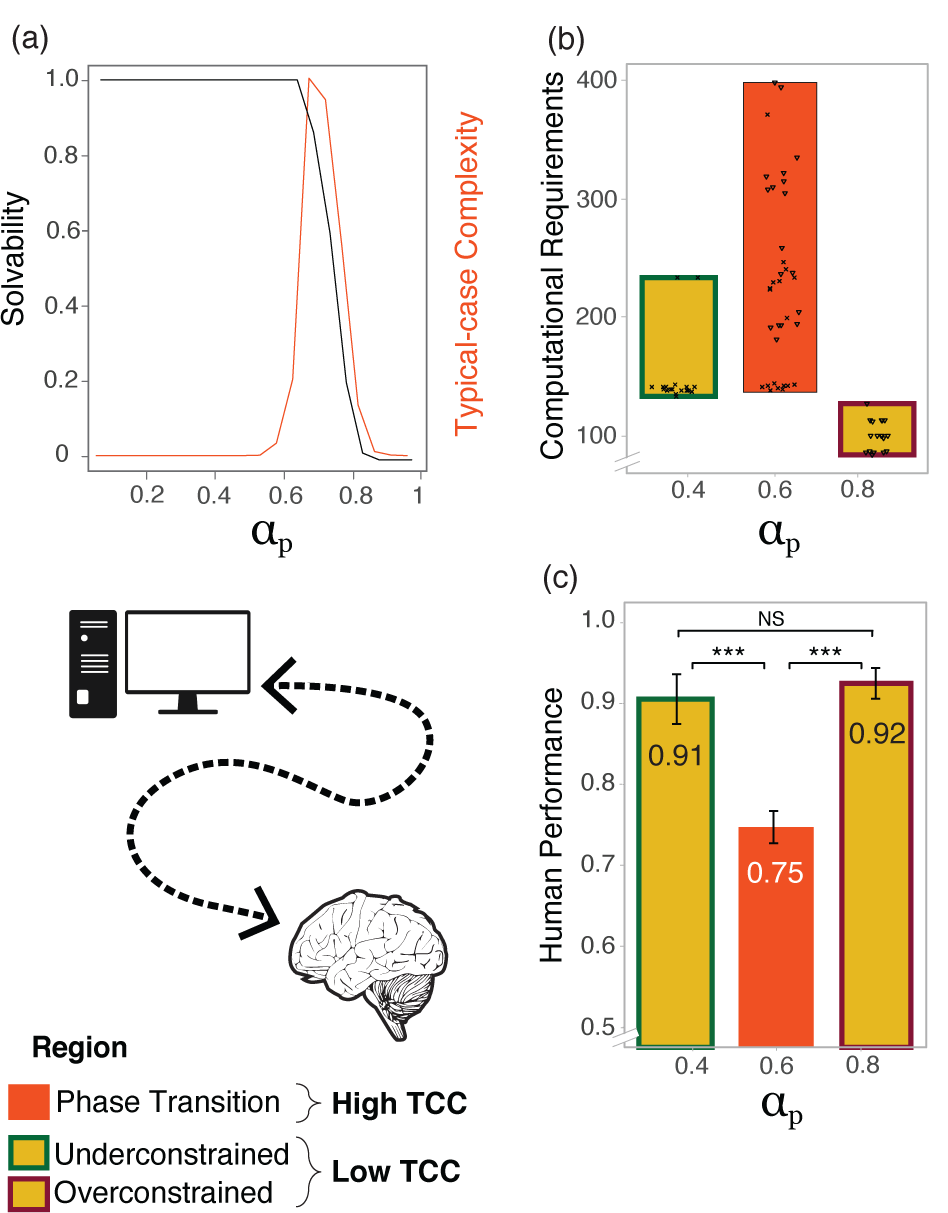
Typical-case Complexity and performance in the knapsack decision task. **(a) Computer performance and the phase transition.** Probability of an instance being *satisfiable* as a function of *α*_*p*_ (left axis). The values presented correspond to the knapsack decision problem with 30 items and **fixed *α***_***c***_ ***≈* 0**.**44**. The right axis shows a pictorial representation of the typical-case complexity (TCC) (28). **(b) Instance sampling for the behavioral experiment**. Each point is an instance sampled as a function of the proxy for computational requirements (number of propagations using the *Gecode* solver) and the normalized profit *α*_*p*_. **(c) Human performance by typical-case complexity in the knapsack decision task**. Mean performance and standard errors. The values presented in (b) and (c) correspond to the knapsack decision problem with 6 items and **fixed *α***_***c***_ **∈ [0**.**40 *−* 0**.**45]** (see Materials and Methods). *Note:* ^*\**^*p<0*.*1;* ^*\*\**^*p<0*.*05;* ^*\*\*\**^*p<0*.*01; NS: not significant*.

In a second approach, we construct a metric of complexity of instances of the decision variant of the knapsack problem that is specific to a single instance. The metric, referred to as instance complexity (IC), is based on a comparison between an instance of the decision variant and the solution of the corresponding instance in the optimization variant. Deriving this metric is more computationally expensive than TCC since it requires computing the solution of the optimization variant. But, unlike TCC, it obviates the need to commit to an ensemble of instances and a distribution over this ensemble.

## Results

We studied how a set of mathematical properties of ensembles of instances (TCC) as well as of individual instances (IC) affect human decision quality and time-on-task in the knapsack problem. Participants first solved 72 instances of the decision variant of the knapsack problem (Fig 2a), followed by 18 instances of the optimization problem (Fig 2b).

**Fig. 2.**
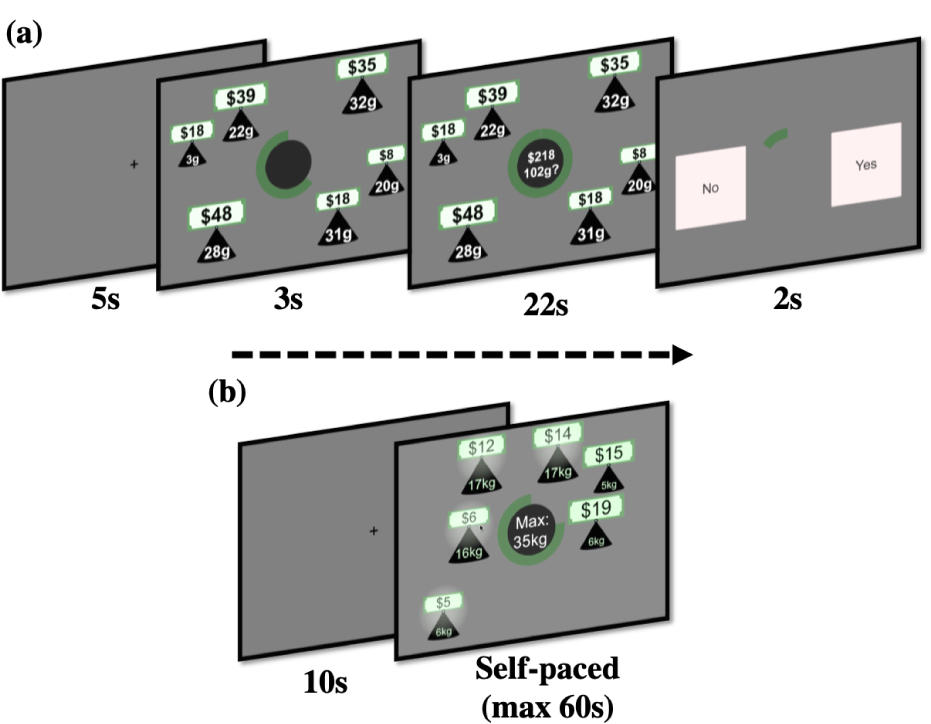
Knapsack Tasks. **(a)** Knapsack decision task. Initially, participants were presented with a set of items of different values and weights. The green circle at the center of the screen indicated the time remaining in this stage of the trial. This stage lasted 3 seconds. Then, both capacity constraint and target profit were shown at the center of the screen. Participants had to decide whether there exists a subset of items for which (1) the sum of weights is lower or equal to the capacity constraint and (2) the sum of values yields at least the target profit. This stage lasted 22 seconds. Finally, participants had 2 seconds to make either a ‘YES’ or ‘NO’ response using the keyboard. A fixation cross was shown during the inter-trial interval (5 seconds). **(b)** Knapsack optimization task. Participants were presented with a set of items of different values and weights together with a capacity constraint shown at the center of the screen. The green circle at the center of the screen indicated the time remaining in this stage of the trial. Participants had to find the subset of items with the highest total value subject to the capacity constraint. This stage lasted 60 seconds. Participants selected items by clicking on them and had the option of submitting their solution before the time limit was reached. After the time limit was reached or they submitted their solution, a fixation cross was shown for 10 seconds before the next trial started.

### Knapsack decision task

#### Summary statistics

All instances in the experiment had *n* = 6 items. The number of items was selected, based on pilot data, to ensure that the task was neither too difficult nor too easy. Mean *human performance*, measured as the percentage of trials in which a correct response was made, was 83.1% (min = 0.56, max = 0.9, *SD* = 0.08).

On average, participants chose the ‘YES’ option in 48.1% of trials (min = 0.32, max = 0.60, *SD* = 0.06). Performance did not vary during the course of the task (*P* = 0.196, main effect of trial number on performance, generalized logistic mixed model (GLMM); S1 Table Model 1), suggesting that neither experience with the task nor mental fatigue affected task performance.

#### The effect of typical-case complexity on performance

Recent work has studied the typical-case complexity (TCC) of random instances of the 0-1 knapsack problem (28). It has been demonstrated that the hardest instances tend to appear in the vicinity of the so-called *satisfiability threshold*, where a phase transition occurs in the probability of an instance being *satisfiable* (that is, the probability that the correct answer to the instance is ‘yes’). In the phase transition, this probability changes precipitously from one to zero. The probability that the instance is satisfiable can be expressed in terms of a small set of instance parameters 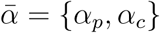:

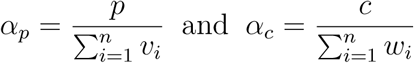

where *w*_*i*_ are the weights of the items, *v*_*i*_ are values of the items, *n* is the number of items, *c* is the weight capacity and *p* is the target profit.

Therefore, there exists a mapping from instance properties to computational complexity of the instance. The phase transition separates instances of the problem into two regions: an under-constrained region where the constraints are lenient, and thus many solutions are likely to exist, and an over-constrained region where the constraints are stringent, and thus the existence of a solution is unlikely (that is, an instance is not satisfiable). Computing the solution of instances in the proximity of the satisfiability threshold requires on average more computational resources than for instances further away from it (Fig 1a), similar to what has been observed in relation to a number of other NP-hard problems (12, 22, 28–31).

The instances in this study were chosen such that they were located at different distances from the satisfiability threshold (Fig 1a). Instances near the phase transition are categorized as having high typical-case complexity (*high TCC*) whereas instances further away from it - that is, in the under-constrained (*α*_*p*_ ∈ [0.35, 0.4]) and over-constrained (*α*_*p*_ ∈ [0.85, 0.9]) regions - are categorized as having low typical-case complexity (*low TCC*). More detail is provided in Materials and Methods, and in S3 Appendix.

In order to test whether participants’ ability to solve an instance was affected by TCC, we first compared performance on instances with high TCC and low TCC. We expected our participants to perform better on instances with low TCC compared to instances with high TCC. Performance was significantly lower on instances with high TCC (*P <* 0.001, main effect of TCC on performance, GLMM; Fig 1c; S1 Table Model 2). This suggests that TCC affected participants’ ability to solve an instance.

To examine this result in more detail, we hypothesized that performance is affected by the tightness of the profit and capacity constraints. We tested whether performance on instances in the under-constrained region (*α*_*p*_ *≈* 0.4) was different to performance on instances in the over-constrained region (*α*_*p*_ *≈* 0.9). We found no significant difference in performance between the two regions with low TCC (*P* = 0.355, main effect of region, GLMM; S1 Table Model 7; Fig 1c), but confirmed a significant difference in performance between instances with high TCC and each of the other two regions (*P <* 0.001, difference in performance between regions, GLMM; S1 Table Model 6).

We also hypothesized that the effect of TCC on performance is affected by the satisfiability of an instance, that is, whether the answer to the decision problem is ‘yes’ or ‘no’. This hypothesis is based on an asymmetry of NP problems. Proving that an instance is *satisfiable* requires finding one subset of items that satisfy the constraints. Such a set may be identified without exploring the full search space and, additionally, there may be more than one such subset. In contrast, to conclude that an instance is *unsatisfiable* requires proving that no such set exists. This might require a full search over every possible subset of items in order to determine that none of the subsets satisfies the constraints. We investigated the effect of satisfiability on performance and found that the effect of TCC was still significant when controlling for satisfiability (*P <* 0.001, main effect of TCC on performance, GLMM; S1 Table Model 3), but that there was no significant effect of satisfiability on performance (*P* = 0.355 main effect of satisfiability on performance, *P* = 0.796 interaction effect of TCC and satisfiability on performance, GLMM; S1 Table Model 3).

#### Structure of an instance and human performance

For satisfiable instances, the tightness of the constraints can be studied further by analyzing the number of subsets of items that satisfy the constraints. We call this the *number of witnesses*. We found that (for satisfiable instances), the probability of solving an instance correctly increases as the number of witnesses (combinations of items that satisfy the constraints) increase (*P* = 0.001, main effect of number of witnesses on performance, GLMM; S1 Table Model 8; Fig 3c). This suggests that participants were more likely to solve an instance correctly when there were more witnesses. Moreover, the probability of solving the instance correctly increased faster (with respect to number of witnesses) if the instance had high TCC (*P <* 0.001, interaction effect of TCC and number of witnesses on performance; GLMM; S1 Table Model 8).

**Fig. 3.**
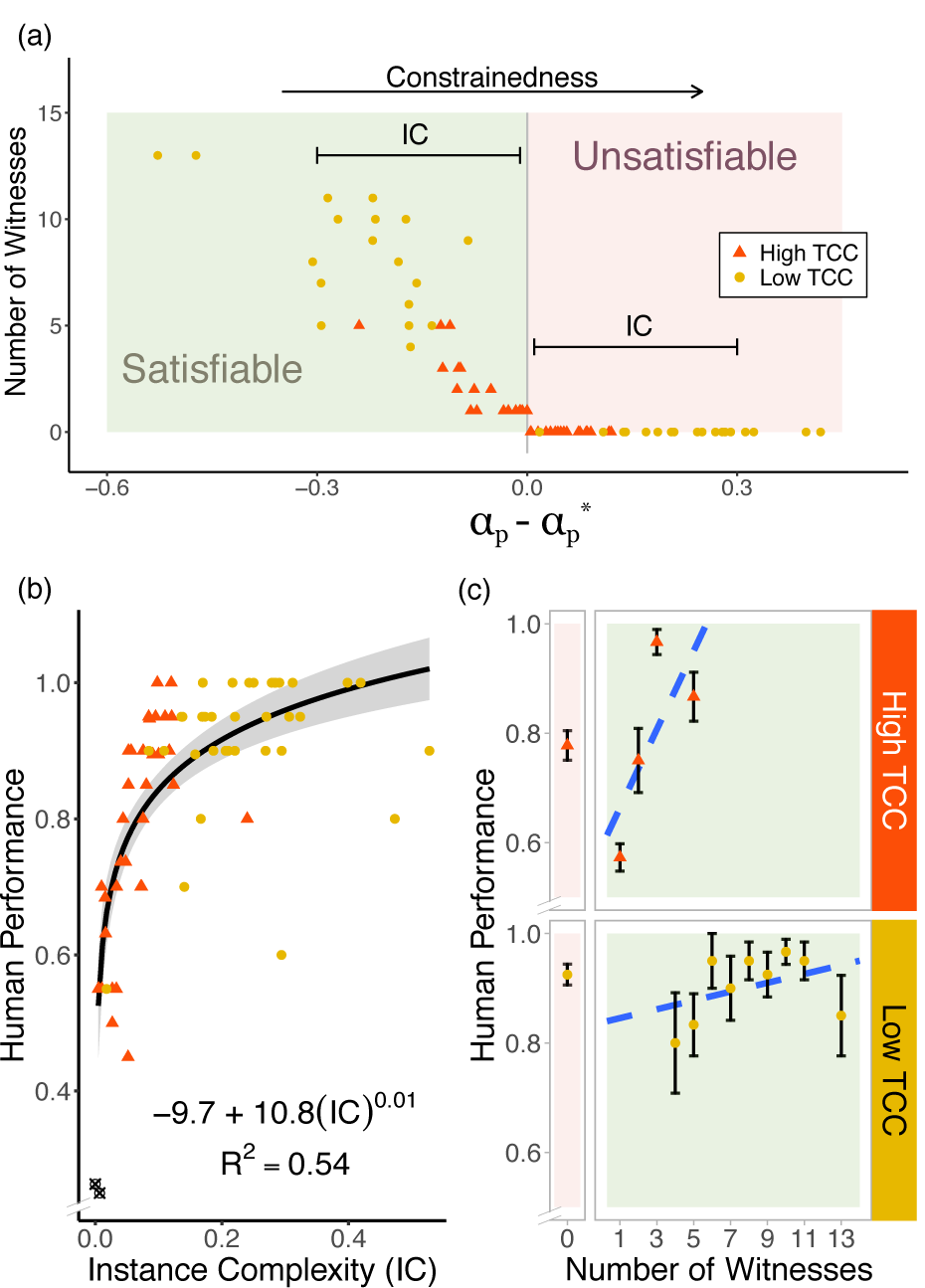
Properties of sampled instances and human performance. **(a)** Properties of sampled instances. Each decision instance is plotted according to how large is the gap between the normalized profit *α*_*p*_ and the maximum achievable normalized profit 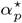 given the set of items and the capacity constraint *c*. Instances become more constrained as 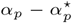 increases. Instances in the negative quadrant are satisfiable and, thus, have at least one solution witness. The number of witnesses is defined as the number of item combinations that satisfy both capacity and profit constraints. Instances in the positive quadrant are all unsatisfiable with 0 witnesses. **(b) Relation between IC and human performance in the Knapsack Decision Problem**. Mean performance and IC by instance. Instances are divided into high and low TCC. Outliers are denoted in black and excluded from the model fit. **(c) Relation between performance and number of witnesses in the knapsack decision task**. Mean performance and standard error by number of solution witnesses.

We further explored the link between TCC, which is related to the satisfiability probability, and the number of witnesses. We studied the theoretical connection and found that the same parameters 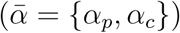 that characterize TCC also describe the expected number of witnesses (see S4 Appendix). We then tested this link empirically and found, as expected, that satisfiable instances with high TCC tend to have a lower number of witnesses than satisfiable instances with a low TCC (*P <* 0.001, unpaired t-test; Fig 3a). Taken together, these findings corroborate that 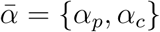 capture the constrainedness of an instance (see S4 Appendix).

#### Instance complexity and human performance

Next, we propose a measure of complexity that is based on the properties of a single instance. We refer to this measure as *instance complexity* (IC). IC is more comprehensive than the number of witnesses (it can quantify complexity for both satisfiable and unsatisfiable instances), and is easier to compute. It is defined based on the distance between the level of the profit constraint (target profit) and the maximum value attainable in the corresponding instance in the optimization variant of the 0-1 knapsack problem, that is, an instance with the same set of items and the same capacity constraint (Fig 3a). Specifically,

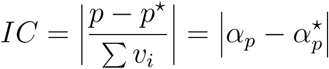

where *p* is the target profit of the decision instance and *p\** is the maximum value achievable of the optimization instance with the same weights and values of items as well as the same capacity constraint. *α*_*p*_ and 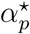 are the corresponding normalized values. Note that a lower IC implies higher computational complexity.

We first explored the relation between IC and TCC. Both measures map constrainedness to computational complexity. However, they do this at different levels. TCC maps average constrainedness of a random ensemble of instances to their average complexity, whereas IC maps the constrainedness of a single instance to its complexity, regardless of which ensemble it was sampled from. Given that both measures map constrainedness to complexity, we expected both measures to be highly correlated. As predicted, instances in our study with *low TCC* had a higher average IC than instances with *high TCC* (*P <* 0.001, *β*_*T CC*_ = *−*0.175, Observations= 70; Fig 3b).

We explored the relation between IC and average human performance on individual instances in the knapsack decision task. We found a positive non-linear relation between this measure and average accuracy (*R*^2^ = 0.542; best *AIC* among competing models; S7 Table; Fig 3b). We also compared the model fit with respect to a model with only TCC as explanatory variable. As would be expected, IC models performs better than the TCC model (*R*^2^ = 0.21; *AIC*_*T CC*_ is highest among competing models; S7 Table). Finally, the effect of IC on performance was further corroborated using a mixed effects model (*P <* 0.001, main effect of *IC*^0.01^ on performance, GLMM; S1 Table Model 9).

#### Algorithm-specific complexity measures and performance

So far, we have analyzed human performance in relation to mathematical properties 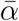 of instances, independent of any particular algorithm or strategy for solving the instance. We now turn to a more fine-grained analysis of computational resource requirements. To do this, we tested whether human performance was related to the number of computational operations of particular (canonical) algorithms.

We considered two widely-known, generic solvers for constraint satisfaction problems, *Gecode* (32) and *Minisat*^*+*^ (33, 34). For each of these solvers, we chose a metric that indicates the difficulty for the algorithm of finding a solution and whose value is highly correlated with compute time. Both metrics quantify the extent of the search effort the respective solver had to undertake to find the solution, which is related to the number of computational steps performed and thus to compute time (see S2 Appendix). We found that human performance in the instances was negatively related to the computational steps used by the Gecode algorithm (*P <* 0.001, main effect of number of propagations, GLMM; S1 Table Model 4). On the other hand, the relation between human performance and the number of computational steps the Minisat^+^ algorithm required was not significant (*P* = 0.395, main effect of number of decisions, GLMM; S1 Table Model 5).

### Knapsack optimization task

#### Summary statistics

We first analyzed participants’ ability to find the optimal solution of an instance. We define *computational performance* as a dichotomous variable that is equal to 1 if the participant obtained a value equal to the maximum value obtainable in the instance, and 0 otherwise. Mean computational performance was 83.2% (min = 0.67, max = 0.94, *SD* = 0.08). Participants spent 43.5 seconds on average on an instance (min = 27.4, max = 60.0, *SD* = 8.9). Participants were allowed to select any set of items, irrespective of the capacity constraint, which implied that they could submit candidate solutions that exceeded the capacity constraint. However, the capacity constraint was only violated in 3% of instances. Performance did not change throughout the task (*P* = 0.683, main effect of trial number on performance, GLMM; S2 Table Model 1), nor did the time spent per instance (*P* = 0.483, main effect of trial number on time, linear mixed model (LMM); S3 Table Model 1), suggesting that neither experience nor mental fatigue affected task performance.

#### Effect of typical-case complexity on performance

We define typical-case complexity for optimization problems (*TCC*_*O*_) as the TCC of the decision problem of choosing whether the optimal value is achievable (see S1 Appendix). We hypothesized that computational performance in instances with *high TCC*_*O*_ (instances whose solutions have a corresponding decision problem with high TCC) would be lower than in instances with *low TCC*_*O*_ (instances whose solutions have a corresponding decision problem with low TCC). We found that mean computational performance was lower in those instances with high TCC_*O*_, relative to those with low TCC_*O*_ (*P <* 0.001, main effect of TCC_*O*_, GLMM; Fig 4a; S2 Table Model 2).

**Fig. 4.**
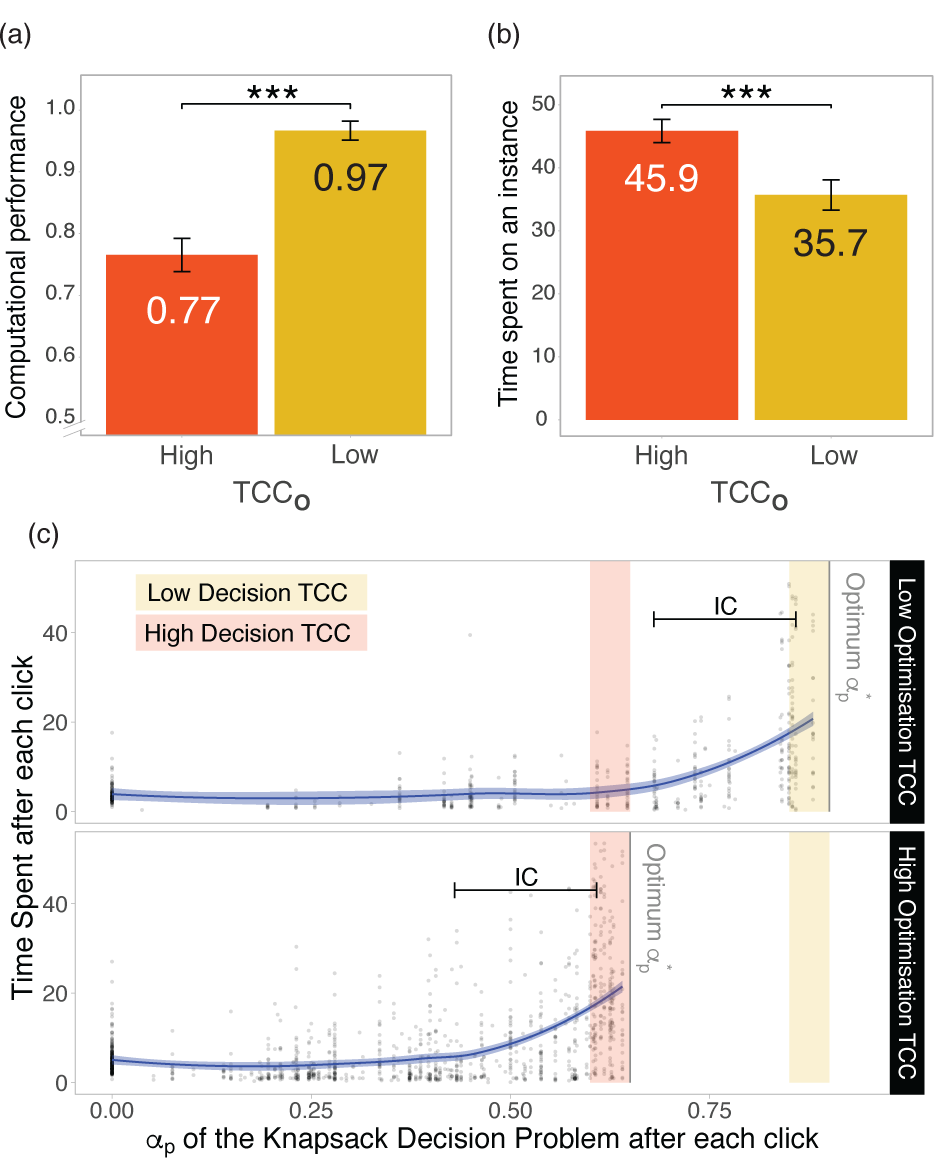
Relation between computational complexity and human performance in the knapsack optimization task. **(a) Relation between TCC**_*O*_ **and computational performance**. Mean computational performance and standard error of the means (SEM) in the knapsack optimization task according to TCC_*O*_. **(b) Relation between TCC**_*O*_ **and time-on-task on an instance**. Mean time spent (and SEM) in the knapsack optimization task according to TCC_*O*_. **(c) Time spent after each click and TCC**. After each click participants were faced with the question: “Is there another set of items with a higher profit that still satisfies the weight capacity constraint?”. Each of these decisions is a Knapsack Decision Problem with a corresponding *α*_*p*_ and corresponding TCC. The figure shows the amount of time people spent at each of these decisions before doing another click. Note that the IC of the Knapsack Decision Problem at each click is defined as the distance between *α*_*p*_ and the optimum 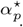. The top panel shows instances with low TCC_*O*_ ; that is, optimization instances whose optimum lies in a low TCC region. The bottom panel shows optimization instances with a high TCC_*O*_. *Note:* ^*\**^*p<0*.*1;* ^*\*\**^*p<0*.*05;* ^*\*\*\**^*p<0*.*01; NS: not significant*.

So far, we have defined computational performance as a dichotomous variable. We now look at a finer-grained measure. To this end, we define *item performance* as the minimum number of item replacements needed to reach the optimal solution. These include both the removal of items that are not in the optimal solution and the addition of items that are in the optimal solution (but not part of the candidate solution). The higher the value of this measure, the further away the submitted solution is from the optimum in item space. We found that item performance was worse, on average, in instances with high TCC_*O*_, relative to instances with low TCC_*O*_ (*P <* 0.001, main effect of TCC_*O*_, LMM; S4 Table Model 2).

Another way of defining performance is in terms of value obtained in an instance. We define *economic performance* as the ratio of the total value of items in the submitted solution to the total value of items in the optimal solution. We found that economic performance was lower in instances with high TCC_*O*_ relative to instances with low TCC_*O*_ (*P <* 0.001, main effect of TCC_*O*_, LMM; S4 Table Model 1).

#### Relation between performance in the knapsack decision task and the knapsack optimization task

We hypothesized that participants’ performance in the two tasks is related and that participants who performed better in the knapsack decision task performed better in the knapsack optimization task. For this analysis we excluded one participant with performance in the knapsack decision task significantly below the performance of any other participant. We found a positive and significant correlation between performance in the two tasks (Pearson correlation = 0.67, *P* = 0.002, d.f. = 17, correlation between average performance in the decision variant and computational performance in the optimization variant). If the outlier is included, we get qualitatively similar results (Pearson Correlation = 0.49, *P* = 0.027, d.f. = 18).

#### Relation between complexity and time-on-task

The knapsack optimization task also allowed us to investigate a relevant aspect of effort, namely time-on-task. As we did not incorporate any direct opportunity costs of time in our experimental setting, clock time does not capture this aspect of effort. However, clock time increases in the number of computations performed, as well as the time required for each computation.

We hypothesized that participants would expend more time on more difficult instances. As expected, participants spent more time on instances with high TCC_*O*_ relative to those with low TCC_*O*_ (*P <* 0.001, main effect of TCC_*O*_, LMM; Fig 4b; S3 Table Model 2). This effect was also present when controlling for computational performance (*P* = 0.037, main effect of TCC_*O*_, LMM; S3 Table Model 6). This means that even when participants did not find the optimal solution, they expended more time on instances with high TCC_*O*_.

Next, we analyzed the relation between time expended in an instance and performance in the instance. We found a negative relation between time-on-task and the probability of finding the solution (*P <* 0.001, main effect of time, GLMM; S2 Table Model 7). However, when we account for TCC_*O*_, the effect of time on performance is no longer significant (*P* = 0.905, main effect of time; *P* = 0.352, interaction effect of time and TCC_*O*_, GLMM; S2 Table Model 3). Taken together with previous results, it appears that the relation between time-on-task and computational performance is driven by TCC_*O*_. That is, the negative correlation between time expended and performance may have been caused by the separate effects of TCC_*O*_ on time expended and performance.

In order to further examine the relation between optimization instances, time expended and complexity, we examined the amount of time participants spent after each click at each selection of items before performing the next click. After each click participants were faced with the question: “Is there another set of items with a higher profit that still satisfies the weight capacity constraint?”. Previous results suggest that TCC of the decision problem faced after each click has an effect on the time spent at each selection of items (28). We explored this link and found that the effect was driven by the constrainedness of the instance (Fig 4c). We found that the time spent after each click was mainly influenced by IC (*P <* 0.001, main effect of IC, LMM; S5 Table Model 2) rather than by TCC (*P* = 0.206, main effect of TCC, LMM; S5 Table Model 2). Additionally, at each click, we estimated how many subsets of items would yield a greater sum of values than the current selection, while still satisfying the capacity constraint. We found that participants spent more time at those stages where there were fewer alternatives that yielded a more valuable solution, whilst still satisfying the capacity constraint (*P <* 0.001, main effect of the number of more valuable solutions, LMM; S5 Table Model 1).

#### Relation between algorithm-specific complexity measures, time-on-task and performance

We next examined a set of alternative complexity measures based on the generic solution algorithms *Gecode* and *Minisat*^*+*^. We found qualitatively similar results to those for the decision variant of the knapsack problem, with higher instance difficulty, according to *Gecode* propagations associated with lower average performance (*P <* 0.001, main effect of number of propagations, GLMM; S2 Table Model 4). For the *Minisat*^*+*^ number of decisions this effect was not significant (*P* = 0.157, main effect of number of decisions, GLMM; S2 Table Model 5).

We also examined whether these complexity measures were related to the time spent on each of the instances. We found that instances with a higher number of Gecode propagations were associated with higher time-on-task (*P <* 0.001, main effect of number of propagations, LMM; S3 Table Model 3). We found a similar relation for the Minisat^+^ decision measure (*P* = 0.001, main effect of number of decisions, LMM; S3 Table Model 4), which captures the positive relation between time expended and the computational requirements of solving an instance using the algorithm.

We also analyzed the relation between computational performance and *Sahni-k*, another algorithm-specific measure of complexity. *Sahni-k* is proportional to both the number of computations and the amount of memory required to solve an instance of the knapsack optimization task. In general, a higher *Sahni-k* is related to higher computational resource requirements. This metric has previously been shown to be associated with performance in the knapsack optimization task (4, 18). In line with these studies, we found a negative relation between *Sahni-k* and computational performance (*P <* 0.001, main effect of *Sahni-k*, GLMM; S2 Table Model 6) and a positive relation between *Sahni-k* and time-on-task (*P* = 0.001, main effect of *Sahni-k*, LMM; S3 Table Model 5). However, when controlling for TCC, the effect of *Sahni-k* on time-on-task is no longer significant (*P* = 0.580, main effect of *Sahni-k*, LMM; S3 Table Model 7).

### Relation between performance in knapsack tasks and cognitive function

Finally, we investigated the relation between performance in the two knapsack tasks and various aspects of cognitive function. In particular, we used tests aimed at assessing mental arithmetic, working memory, episodic memory, strategy use as well as processing and psycho-motor speed. Correlations between performance in these tasks and the knapsack tasks were all non-significant (see Materials and Methodsand S6 Table for details).

## Discussion

In this study, we examined the effect of intrinsic complexity of instances on both performance and time-on-task in both the decision variant and the optimization variant of the 0-1 knapsack problem, both NP-hard tasks. To this end, we used two measures of computational hardness that are based on mathematical properties of instances. The first, typical-case complexity (TCC), captures the average complexity of random ensembles of instances. The second, instance complexity (IC), is a metric of complexity of individual instances.

We tested the effect of both measures of complexity on human problem-solving. We found that performance was lower in random instances with high TCC in both variants of the knapsack problem compared to instances with low TCC. Moreover, time expended was positively correlated with TCC. We synthesized the features that characterize TCC in a new measure of instance complexity (IC) and found a strong effect of this measure on time-on-task and performance. Thus, we provide evidence that human ability to solve complex computational problems is related to mathematical properties of individual instances of such problems.

Many models of decision-making, implicitly or explicitly, require the decision-maker to solve computationally intractable problems, that is, problems that are NP-hard (1, 2). However, little is known about the generic effect of computational complexity on human decision-making. Following a popular approach to studying computational complexity of NP-hard problems (11, 12, 21, 22, 28, 30, 31, 35), we study which properties of instances of those problems drive extreme hardness for humans.

This study is the first to investigate the relation between generic structural properties of individual instances of an NP-hard computational problem and human behavior. We explore, in particular, the effect of the satisfiability and constrainedness on human performance. Our results provide strong evidence that properties of individual instances of computationally hard problems determine extreme hardness for both human and digital computers. This can be interpreted as further evidence that computational complexity is an inherent (or absolute) property of instances of such computational problems (3, 11, 12).

### Computational complexity and models of decision-making

It has been suggested that models of decision-making, and cognition more generally, ought to be computationally tractable in order to be plausible from a computational point of view (2, 36–40). The P-Cognition Thesis proposes that computational plausibility should be linked to P-time computability (36, 39). However, this thesis is often considered too restricted (37, 40). Recognizing that instances of NP-hard problems can vary substantially in computational resource requirements, the Fixed Parameter Tractable (FPT) Cognition Thesis requires models be computable in polynomial time at least for some fixed input parameter (1, 37).

Our study both supports these proposals but also complements them. It provides empirical evidence relating human decision-making capacity to individual properties of instances and shows which of those properties make individual instances hard for people. Thus, it supports the assumption of frameworks like the P-Cognition Thesis (2, 36–39) that computational complexity affects human decision-making capabilities and that many existing models of decision-making cannot possibly be generic given that their implementation may require computational capacities beyond those available to people. Additionally, our study demonstrates that the study of hardness of individual instances and of random ensembles provides insights that can be used to predict both human performance and time-on-task. This approach could be used to refine frameworks like fixed parameter tractability (1, 37). Specifically, it could be used to explore how the hardness of particular instances of a problem are related to worst-case asymptotic complexity.

Moreover, our results have a significant implication in the approach used to study human decision-making. Previous findings suggest that humans deploy a vast range of strategies (4, 41, 42), thus limiting the predictive power of current models of decision-making. Here, we postulate that performance and time-on-task are driven by properties of the instance at hand rather than being an idiosyncratic feature of the solver (the human), and suggesting that predictive power of decision theoretic models can be improved by including instance properties. That is, the goal so far has been to identify the procedures (algorithms, heuristics) that humans deploy in search of a solution (41, 43), while instead performance and time-on-task may be more readily predicted from properties of an instance.

### Difficulty of cognitive tasks and computational complexity

It has previously been suggested that computational complexity could be used to measure difficulty of cognitive tasks. We would argue that this is indeed the case but that complexity classes such as P or NP are too broad for this purpose given that they are based on asymptotic worst-case approaches. We propose that TCC and IC are better suited to study difficulty of tasks for humans.

TCC is a more fine-grained measure that studies complexity of “typical” instances of a problem. Moreover, it captures complexity in a way that is independent of a particular algorithm or model of computation (11, 12, 30) and it has been proven to be applicable to a large range of problems, including the graph coloring problem (12, 22), the traveling salesperson problem (29) and the K-SAT problems (Boolean satisfiability problems) (12, 22, 30, 31). Our findings show that TCC has an effect on performance, as well as on time-on-task.

We also investigated IC as an alternative metric to capture difficulty of an instance and we find a close relation between this measure and human performance. IC complements TCC by providing a measure of difficulty at an individual instance level. While TCC maps the average constrainedness of an ensemble of random instances to its average complexity, IC maps the constrainedness of a single sampled instance of that random ensemble (after the instance has been generated) to its complexity. In other words, IC measures difficulty of a single instance and it does not depend on a random instance generation process. It is worth noting, however, that TCC has a key advantage over IC. In order to compute IC, the corresponding optimization problem needs to be solved, whereas TCC is a measure that can be estimated entirely from mathematical properties of the problem. This makes TCC not only less computationally intensive, but perhaps also a better candidate for playing a role in human meta-decisions such as strategy selection (44, 45).

This combination of findings provides support for the conceptual premise that computational complexity constitutes a core aspect of task difficulty. Here, we looked at time complexity. We would point out that time complexity is likely not the only measure that captures difficulty of tasks for participants. There may be many other relevant cognitive dimensions, such as memory, that are relevant for understanding cognitive limitations (40, 46, 47).

In order to explore other dimensions of difficulty, we tested participants on a set of cognitive abilities including attention, working memory and mental arithmetic. Individual differences in performance in these tasks were independent of individual differences in performance in the knapsack tasks. One possible explanation for the lack of correlation is that these cognitive abilities play only a minor role in solving computationally hard problems and that those problems instead require another cognitive ability that is not captured by any of the tests we administered. Another possible explanation is that we did not measure the active cognitive constraints that drove differences in individual performance. It is, of course, also possible that our study did not have sufficient statistical power to detect individual differences. Further research is needed in order to incorporate the full spectrum of cognitive costs and resource limitations and link them to performance and time-on-task in decision tasks.

### Meta-decision making and adaptation of strategies

In order for a theory of decision-making to be plausible from a computational perspective, the computational requirements of a decision task need to be within the cognitive resources available to a decision-maker. Indeed, it has been suggested that the principle of rationality should not (only) be applied at the level of behavior (Marr’s computational level) but (also) at the level of computation (Marr’s algorithmic level), an approach known as resource rationality (48). In this framework, limited computational resources are allocated to tasks in a way that is optimal relative to specified objective function (48). In order to understand how limited computational resources are allocated, it is necessary to study both the cognitive capacities of decision-makers as well as the cognitive requirements of a task. Our study provides a framework for studying the interaction between the two; that is, the computational limitations in a task. Specifically, they can be measured using insights from computational complexity theory to predict performance and compute time.

A relevant dimension of expected costs of performing a task is, arguably, the computational resource requirements of performing the task. Empirically, evidence from the current study suggests that agents expend more time on instances with higher TCC. In particular, we found that that the time participants spent in the task was modulated by TCC. Theoretically, TCC is particularly suitable as an approximation of the expected computational requirements because of its characteristics. Firstly, it is an *ex-ante* measure, that is, it is based on a set of features of the task, which could potentially be identified and used by the agent *before* solving the task. Secondly, the set of features related to TCC are intrinsic to the task, that is, they are not specific to the particular algorithm used to solve a problem. Thirdly, TCC has been shown to be generalizable to a large set of computational problems (12, 28–31). Further research could usefully explore whether humans compute TCC in order to estimate the expected costs of performing a task.

Our approach provides a framework that lends itself to studying why particular heuristics or algorithms are successful on some instances but not on others. Moreover, it could explain why participants’ use of heuristics changes with instance properties (49, 50). It is worth noting, however, that the problem of choosing a heuristic among a set of possible heuristics can in itself be an NP-hard problem (51). Further research could explore how TCC affects strategy selection, and in particular heuristic selection.

### Directions for future research

We have studied an important aspect of difficulty, for humans, of solving random instances of the knapsack problem, which is ubiquitous in daily life (15, 18, 52). However, there are many other problems that are relevant for humans. The framework of TCC has been shown to generalize to other NP-hard problems (12, 28–31). It is an open question whether the applicability of TCC and IC to human problem-solving extends to these other problems as well. Future work should address this question.

Furthermore, in our study, the optimization task involved finding the optimal solution. However, finding the exact solution might not always be required in the real-world. In many cases, finding an approximate solution might suffice. Future research should investigate whether the results found in this study, for humans and other types of computers, can be extended to approximation instances. Nevertheless, it is worth noting that for some hard problems, approximating their solution can be computationally as hard as computing the solution itself (3, 53).

Our results for TCC are based on a particular sampling distribution. Specifically, we used the uniform distribution to sample the knapsack instances. This approach has been used to understand hardness and to study “typical” instances of a problem, but these instances might not necessarily be the ones we encounter outside of the laboratory setting. Characterizing real-life distributions of instances is an open research question in computer science (54). Further research would be required to study whether this method is generalizable to other sampling distributions and, specially, to those distributions that are encountered in everyday life.

Our work provides a step towards understanding the effects of computational complexity on human behavior by providing a measure of expected decision difficulty. We have shown that TCC and IC affect behavior through task performance. Yet, it could also have an impact on behavior in other ways. For instance, attitudes towards complexity could affect behavior. Complexity avoidance could lead people to avoid situations that involve solving difficult tasks, whereas complexity seeking could lead to situations in which people seek tasks that require a high amount of effort to be solved (55). Another way that complexity could be related to behavior is through its effect on uncertainty. In the case of the knapsack optimization task, it is still an open question which mechanism participants used to adjust the time-on-task. TCC could influence the level of uncertainty of having found the solution, and in turn this uncertainty could play a role in the decision of when to submit an answer (56). We leave it to future work to explore the effects of attitudes towards (or preferences over) complexity in decision-making, as well as the relation between complexity, uncertainty and behavior. This study provides evidence that computational complexity is an inherent property of a computational problem and, in particular, of an instance (57). This supports the thesis that computational complexity affects problem solving across computing systems, such as von Neumann architectures and the brain, thus supporting frameworks that use computational complexity to study cognition, such as the FPT Cognition Thesis (37).

In a broader context, the present study might help to identify the limits of human cognition and decision-making; thus, providing a building block for the development of a theory of complexity of human computation (40). This is crucial for the design of policies that wish to improve the quality of decisions people make and the outcomes they achieve in areas such as financial investments or the selection of health insurance contracts, among many others. In those cases where the task is too demanding, mechanisms could be designed to help people improve the quality of their decisions. This could be done, for instance, through software applications that take advantage of the computational power of electronic computers. Finally, our results advocate for closer collaboration between decision scientists and computer scientists. Not only can decision sciences be informed by computation theory, as was done in this study, but research on humans could motivate the development of new theories and algorithms.

## Materials and Methods

### Ethics statement

The experimental protocol was approved by the University of Melbourne Human Research Ethics Committee (Ethics ID 1749616). Written informed consent was obtained from all participants prior to commencement of the experimental sessions. Experiments were performed in accordance with all relevant guidelines and regulations.

### Participants

Twenty human volunteers recruited from the general population took part in the study (14 female, 6 male; age range = 18-31 years, mean age = 22.0 years). Inclusion was based on age (minimum = 18 years, maximum = 40 years).

### Knapsack decision task

In this task, participants were asked to solve a number of instances of the (0-1) knapsack decision problem. In each trial, they were shown a set of items with different values and weights as well as a capacity constraint and a target profit. Participants had to decide whether there existed a subset of those items for which (1) the sum of weights is lower or equal to the capacity constraint and (2) the sum of values yields at least the target profit.

Each trial had four stages. In the first stage (3 seconds), only the items were presented. Item values, in dollars, were displayed using dollar bills and weights, in grams, were shown inside a black weight symbol. The larger the value of an item, the larger the dollar bill was in size. Similarly, the larger the weight of an item, the larger its weight symbol was in size. At the center of the screen, a green circle indicated the time remaining in this stage. In the second stage (22 seconds), target profit and capacity constraint were added to the screen inside the green timer circle. In the third stage (2 seconds), participants saw a ‘YES’ or ‘NO’ buttons on the screen, in addition to the timer circle, and made a response using the keyboard (Fig 2a). A fixation cross was then shown (5 seconds) before the start of the next trial.

Each participant completed 72 trials (3 blocks of 24 trials with a rest period of 60 seconds between blocks). Each trial presented a different instance of the knapsack decision problem. The order of instances was randomized for each participant.

We excluded a total of 13 trials (from 8 participants) in which no response was made.

All instances in the experiment had 6 items. Instances varied in their computational complexity. It has been shown that computational complexity of instances in the 0-1 knapsack decision problem can be characterized in terms of a set of instance properties (28). In particular, TCC can be characterized in terms of the normalized capacity constraint *α*_*c*_ (capacity constraint normalized by sum of all items weights) and the normalized target profit *α*_*p*_ (target profit normalized by sum of all item values) (Fig 1a). We made use of this property to select instances for the task (see S3 Appendix; Fig 1b).

We selected the normalized capacity bin of *α*_*c*_ ∈ [0.40, 0.45] and chose the normalized profit bins that corresponded to the under-constrained (*α*_*p*_ ∈ [0.35, 0.4]), phase transition (*α*_*p*_ ∈ [0.6, 0.65]) and over-constrained (*α*_*p*_ ∈ [0.85, 0.9]) regions. We randomly selected 18 instances from the under-constrained bin and 18 from the over-constrained bin. Additionally, we sampled 18 *satisfiable* instances and 18 *unsatisfiable* instances near the satisfiability phase transition (*α*_*p*_ ∈ [0.6, 0.65]). Instances near the phase transition are defined to have a *high TCC*, whereas instances further away from the phase transition (in the under-constrained or over-constrained regions) are defined to have a *low TCC*. Throughout we ensured that no weight/value combinations were sampled twice. In order to also ensure enough variability between instances in the phase transition (high TCC) we added an additional constraint in the sampling. We forced half of the instances from the phase transition to have high computational requirements (top 50%), according to an algorithm-specific ex-post complexity measure of a widely-used algorithm (*Gecode*; see S2 Appendix). Analogously, the other half was selected to have low computational requirements (bottom 50%).

### Knapsack optimization task

In this task, participants were asked to solve a number of instances of the (0-1) knapsack optimization problem. In each trial, they were shown a set of items with different weights and values as well as a capacity constraint. Participants had to find the subset of items that maximizes total value subject to the capacity constraint. This means that while in the knapsack decision problem, participants only needed to determine whether a solution existed, in the knapsack optimization problem, they also needed to determine the nature of the solutions (i.e. the items in the optimal knapsack).

The task had two stages. In the first stage (60 seconds), the items were presented together with the capacity constraint and the timing indicator. Items were presented like in the knapsack decision task. Unlike the decision task, however, participants were able to add and remove items to/from the knapsack by clicking on the items. An item added to the knapsack was indicated by a light around it (Fig 2b).

Participants submitted their solution by pressing the button ‘D’ on the keyboard before the time limit was reached. If participants did not submit within the time limit, the items selected at the end of the trial were automatically submitted as the solution. Participants were then shown a fixation cross (10 seconds) before the start of the next trial.

Each participant completed 18 trials (2 blocks of 9 trials with a rest period of 60 seconds between blocks). Each trial presented a different instance of the knapsack optimization problem. The order of presentation of instances in the task was randomized for each participant.

We excluded 2 trials (from 2 participants) because solutions were submitted after less than 1 second into the task. Additionally, 3 participants were excluded from the analysis of submission times because they never submitted a solution before the time-out. This behavior suggests that these participants might have failed to understand the submission instructions.

The characterization of complexity using TCC is based on the satisfiability probability and, therefore, only directly applicable to decision problems. We propose a way in which TCC can be extended to optimization problems by framing the optimization problem as a sequence of decision problems: “Is there another set of items with a higher profit that still satisfies the weight capacity constraint?”. In other words, we model the decision-maker as selecting a subset of items that satisfy the capacity constraint and then decides whether there exist another combination that would yield them a higher profit and still satisfy the constraint. If the answer is yes, the agent chooses one of such combinations and asks themself the same question again. This process is repeated until the answer is no, which means that the optimum has been reached. We approximate the TCC of an optimization problem to be the TCC of the decision problem after reaching the optimum solution. That is, the decision problem of choosing whether the optimal value is achievable (see S1 Appendix).

To generate instances for the task, a sampling process similar to the one for the knapsack decision task was used (see S3 Appendix for more information). We first selected the same normalized capacity bin as for the knapsack decision task (*α*_*c*_ ∈ [0.4 *−* 0.45]). Afterwards, in order to estimate the normalized profit of the optimization problem, we calculated the optimal set of items *A*^*\**^ *∈ A* for each optimization instance. We then estimated the corresponding optimal sum of values (*p*^*\**^ = ∑_*i∈A*_*\* v*_*i*_). The normalized profit was then calculated by dividing the target profit by the sum of values of all of the items 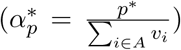.

The normalized profit was then selected to lie in the same regions as in the knapsack decision task and optimization TCC (TCC_*O*_) was defined accordingly. 12 instances were selected from the *high TCC*_*O*_ region 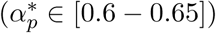 and 6 were selected from the *low TCC*_*O*_ region 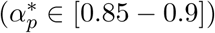. It is worth noting that this process did not generate instances in the under-constrained region 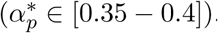.

In order to also ensure enough variability between instances with high TCC_*O*_ we added the same additional constraint in the sampling as in the knapsack decision task. We forced half of the instances with high TCC_*O*_ to have high computational requirements (top 50%), according to an algorithm-specific ex-post complexity measure of a widely-used algorithm (*Gecode*; see S2 Appendix). Analogously, the other half was forced to have low computational requirements (bottom 50%).

### Mental arithmetic task

In this task, participants were presented with 33 mental arithmetic problems (58). The first three trials were considered test trials and thus were not included in the analysis. They were given 13 seconds to solve each problem. The task involved addition and division of numbers, as well as questions in which they were asked to round to the nearest integer the result of an addition or division operation.

### Basic cognitive function tasks

In addition, we also tested participants’ performance on four aspects of cognitive function that we considered relevant for the knapsack tasks, namely, working memory, episodic memory, strategy use, as well as processing and psychomotor speed. To do so, we administered the Reaction Time (RTI), Paired Associates Learning (PAL), Spatial Working Memory (SWM) and Spatial Span (SSP) tasks from the Cambridge Neuropsychological Test Automated Battery (CANTAB) (59).

### Procedure

After reading the plain language statement and providing written informed consent, participants were instructed in each of the tasks and completed a practice session for each task. Participants first solved the CANTAB RTI task, followed by the knapsack decision task. Then they completed the CANTAB RTI task again, followed by the knapsack optimization task. Subsequently, they completed the remaining CANTAB tasks in the following order: PAL, SWM and SSP. Finally, they performed the mental arithmetic task and completed a set of demographic and debriefing questionnaires. Each experimental session lasted around two hours.

Participants received a show-up fee of A$10 and additional monetary compensation based on performance. They earned A$0.7 for each correct answer in the knapsack decision task and A$1 for each correct answer in the knapsack optimization task.

### Statistical analysis

All of the generalized logistic mixed models (GLMM) and linear mixed models (LMM) included random effects on intercept for participants. Their *p*-values were calculated using a two-tailed Wald test. All statistical analyses were done in R and mixed models were estimated using the R package lme4 (60).

### Data and Code Availability

The raw behavioral data, the data analysis code and the computational simulations are all available at the Open Science Framework. The knapsack decision task, knapsack optimization task and mental arithmetic task are also available there (project: https://doi.org/10.17605/OSF.IO/T2JV7).

## ACKNOWLEDGMENTS

This research is supported by a University of Melbourne Graduate Research Scholarship from the Faculty of Business and Economics. Bossaerts acknowledges financial support through a R@MAP Chair from the University of Melbourne.

No competing interests declared.

## Supporting Information Legends

**S1 Appendix** S1 Appendix: Extension of TCC to the knapsack optimization problem.

**S2 Appendix** *Gecode* and *MiniSat*^*+*^ complexity measures.

**S3 Appendix** Instance Sampling.

**S4 Appendix** Expected Number of Witnesses and the constrainedness of the solution space.

**S5 Appendix** CANTAB tasks. Description of the Cambridge Neuropsychological Test Automated Battery (CANTAB®).

**S1 Table** Mixed effects logistic regressions on human performance in the knapsack decision task.

**S2 Table** Mixed effects logistic regressions on computational performance in the knapsack optimization task.

**S3 Table** Mixed effects linear regressions on the time spent on an instance in the knapsack optimization task.

**S4 Table** Mixed effects linear regressions on the other performance measures in the knapsack optimization task.

**S5 Table** Effect of the number of item-subsets that perform better that the current selection of items on time before the next click.

**S6 Table** Pearson correlation between knapsack task performance and cognitive abilities.

**S7 Table** Human accuracy and Complexity in the Knapsack Decision Task. Model fit of alternative models.

## Notes

https://doi.org/10.17605/OSF.IO/T2JV7

### Competing Interest Statement

The authors have declared no competing interest.

### Summary of Updates

Revised Introduction and Discussion. Additional results included.

