## Supplementary Appendices and Tables for "Structural properties of individual instances predict human effort and performance on an NP-Hard problem"

### S1 Appendix: Extension of TCC to the knapsack optimization problem

The characterization of complexity using typical-case complexity (TCC) theory is based on the satisfiability phase transition and therefore is only applicable to decision problems. However, in everyday life we are likely to encounter optimization problems. In order to generalize the TCC measure we frame the knapsack optimization problem (KOP) as a sequence of decision problems in which the question to solve can be reduced to a chain of questions: "Is there another set of items with a higher profit that still satisfies the weight capacity constraint?". In other words, we model the decision-maker as selecting a subset of items that satisfy the capacity constraint and then decides whether there exist another combination that would yield them a higher profit and still satisfy the constraint. If the answer is 'yes', the agent chooses one of such combinations and asks himself the same question again. This process is repeated until the answer is no, which means that the optimum has been reached. In order to incorporate this approach into a TCC measure for optimization problems, we generated a mapping of each KOP instance into a knapsack decision problem (KDP) instance in which the optimum value of the KOP was used as the capacity of the KDP instance (Definition 3). Just as with the TCC of the KDP, we expected performance of human participants to be lower in those instances that map into KDP instances with high TCC.

**Definition 1** *0-1 Knapsack optimization problem (KOP) Let* $n \in \mathbb{N}$, $c \in \mathbb{R}$ *and* $\psi = \{(w_1, v_1), ..., (w_n, v_n)\} \in \mathbb{R}^n \times \mathbb{R}^n$. *The KOP is a mapping* $\mathring{K}_n^{(c)}(\psi) : \mathbb{R}^n \times \mathbb{R}^n \to \mathbb{R}$ *such that:*

$$\mathring{K}_n^{(c)}(\psi) = \max_{\mathbf{x}} \sum_{i=1}^{n} x_i v_i$$
$$\textit{subject to} \sum_{i=1}^{n} x_i w_i \leq c,\ x_i \in \{0, 1\}, i = 1, ..., n.$$

**Definition 2** *Knapsack Decision Problem (KDP)*

*Let* $n \in \mathbb{N}$, $c, p \in \mathbb{R}$ *and* $\psi = \{(w_1, v_1), ..., (w_n, v_n)\} \in \mathbb{R}^n \times \mathbb{R}^n$. *A KDP is a mapping* $K_n^{(c,p)} : \mathbb{R}^n \times \mathbb{R}^n \to \{0, 1\}$ *such that:*

$$\bar{K}_n^{(c,p)}(\psi) = \begin{cases} 1 & \textit{if } \exists A \subseteq \psi \textit{ s.t. } \sum_{(w_i,v_i)\in A} w_i \leq c \textit{ and } \sum_{(w_i,v_i)\in A} v_i \geq p \\ 0 & \textit{otherwise} \end{cases}$$

**Definition 3** *A mapping m from a KOP to a KDP. m is defined as a mapping from a KOP instance* $\mathring{K}(\cdot)$ *to a KDP instance* $\bar{K}'_n(\cdot)$ *such that:*

$$\bar{K}'^{(c,p)}_n(\psi) = \bar{K}_n^{(c,p')}(\psi) \quad \textit{where} \quad p' = \mathring{K}_n^{(c)}(\psi)$$

### S2 Appendix: *Gecode* and *MiniSat*+ complexity measures

We examined a set of alternative ex-post complexity measures based on two existing generic off-the-shelf solvers that are based on different solving algorithms, *Gecode* (1) and *MiniSat*$^+$ (2, 3). Gecode is a constraint solver that uses a constraint propagation technique with different search methods, such as branch-and-bound. We implement this solver using Minizinc (4). Minisat$^+$, on the other hand, transforms the problem into a sequence of satisfiability problems that are then solved using constraint propagation, specifically unit-clause propagation, and backtracking. The corresponding *algorithm-specific* complexity is generally quantified by the time required for the algorithm to solve the problem. However, computational time was not directly used, given that instances with only 6 items are solved rapidly by a computer, and thus the signal-to-noise ratio of this measure is low.

We chose complexity measures that provide a good proxy of the search effort. We tested how well these proxies correlated with computational time of solving knapsack decision and optimization instances with 15, 20, 25 and 30 items. It is worth noting that these complexity measures are not entirely comparable across the two variants of the knapsack problem given that solving different problems might involve different algorithms within one solver. However, we found that the correlations of the complexity measures to computational time are similar across both problems (see Table 1 and 2). We selected those measures with the highest correlation to computational time for the knapsack decision problem (see Table 1). We used these two measures for the knapsack optimization problem as well. For the Gecode solver we used the number of propagations, which is an approximation of the number of options available for exploration after constraint implementation. On the other hand, for the Minisat$^+$ solver we used the number of decisions, which measures the number of assumptions (i.e. number of variable assignments) tried before finding a solution. Both metrics measure the search effort the respective solver had to make to find the solution, which is related to the number of computational steps performed and thus to computational time.

**Table 1. Algorithm-specific complexity measures in the knapsack decision problem. Spearman's correlations between solver time and other solver output variables for the knapsack decision problem. The correlations were calculated for different sizes of the knapsack using over 300,000 simulations for each knapsack size. All correlations are significant at $p < 0.001$.**

**(a) Gecode**

| | Number of Items | | | |
|---|---|---|---|---|
| | 15 | 20 | 25 | 30 |
| **propagations** | **0.95** | **0.99** | **1.00** | **1.00** |
| nodes | 0.95 | 0.98 | 0.99 | 1.00 |
| failures | 0.95 | 0.98 | 0.99 | 1.00 |
| peak_depth | 0.46 | 0.24 | 0.18 | 0.08 |

**(b) MiniSat+**

| | Number of Items | | | |
|---|---|---|---|---|
| | 15 | 20 | 25 | 30 |
| restarts | 0.84 | 0.71 | 0.44 | 0.29 |
| conflicts | 0.96 | 0.86 | 0.97 | 0.99 |
| **decisions** | **0.96** | **0.88** | **0.98** | **0.99** |

**Table 2. Algorithm-specific complexity measures in the knapsack optimization problem. Spearman's correlations between solver time and other solver output variables for the knapsack optimization problem. The correlations were calculated for different sizes of the knapsack using over 2,500 simulations for each knapsack size. All correlations are significant at $p < 0.001$.**

**(a) Gecode**

| | Number of Items | | | |
|---|---|---|---|---|
| | 15 | 20 | 25 | 30 |
| **propagations** | **0.77** | **0.90** | **0.98** | **0.99** |
| nodes | 0.76 | 0.90 | 0.98 | 0.99 |
| failures | 0.76 | 0.90 | 0.98 | 0.99 |
| peak_depth | -0.16 | -0.27 | -0.35 | -0.39 |

**(b) MiniSat+**

| | Number of Items | | | |
|---|---|---|---|---|
| | 15 | 20 | 25 | 30 |
| restarts | 0.81 | 0.87 | 0.92 | 0.96 |
| conflicts | 0.93 | 0.97 | 0.98 | 0.99 |
| **decisions** | **0.81** | **0.90** | **0.96** | **0.98** |

### S3 Appendix: Instance sampling

**Knapsack decision problem.** All instances in the experiment had 6 items. The probability that a particular instance is satisfiable can be characterized in terms of the normalized capacity constraint ($\alpha_c = \frac{c}{\sum_{i=1}^{n} w_i}$) and the normalized target profit ($\alpha_p = \frac{p}{\sum_{i=1}^{n} v_i}$). This probability exhibits a phase transition and the TCC of instances close to the phase transition is higher than further away from the phase transition (see main text).

We made use of this property to select instances for the task, as follows. We first sampled (with replacement) a collection of 250 combinations of weights and values ($< w_1, ..., w_6 >< v_1, ..., v_6 >$) from a uniform (and discrete) distribution over the range

1 to 50. For every weight/value combination, multiple knapsack instances were generated by increasing (in discrete steps) both capacity and the profit constraints from a lower bound of 1 to an upper bound equal up to the sum of weights ($\sum_{i=1}^{n} w_i$) for the capacity constraint ($c$) and the sum of values ($\sum_{i=1}^{n} w_i$) for the target profit ($p$). Target profits and capacities were rounded to the nearest integer and repeated instances were omitted. This process generated a total of 2,496,603 instances. Using these instances, we binned them into bins of width $0.05 \times 0.05$ according to their normalized capacity ($\alpha_c$) and normalized profit ($\alpha_p$). This allowed us to estimate for each one of these bins the probability that the instance was satisfiable.

From the instances generated we selected the normalized capacity ($\alpha_c$) bin of $[0.40, 0.45]$ and chose the normalized profit bins that corresponded to the under-constrained (low TCC; $\alpha_p \in [0.35, 0.4]$), phase transition region (high TCC; $\alpha_p \in [0.6, 0.65]$) and over-constrained (low TCC; $\alpha_p \in [0.85, 0.9]$ regions. We then randomly selected 18 instances from the under-constrained bin and 18 from the over-constrained bin. Finally, we sampled 18 *satisfiable* instances and 18 *non-satisfiable* instances from the phase transition bin (0.4-0.45). Throughout we ensured that no weight/value combinations were sampled twice. In order to also ensure enough variability in instances in the phase transition region we added an additional constraint in the sampling from each bin. We forced half of the instances selected in each bin in the phase transition to be easier than the median according to an algorithm specific ex-post complexity measure (*Gecode* propagations parameter) and the other half to be harder than the median.

**Knapsack optimization problem.** To generate instances for the task, a sampling process similar to the one for the Knapsack Decision Task was used. Taking the same combinations of weights and values sampled for the Knapsack Decision Task, a series of optimization problem instances were generated by increasing (in discrete steps) the capacity constraint from a lower bound of 1 to an upper bound equal to the sum of weights ($\sum_{i=1}^{n} w_i$). Capacities were rounded to the nearest integer and repeated instances were omitted. This resulted in 24,892 instances. To compute the optimization typical-case complexity ($\text{TCC}_O$) of each of these instances, we computed the normalized value of the solution ($\alpha_p^*$ ;see S1 Appendix for details). We selected the same normalized capacity bin as for the Knapsack Decision Task ($\alpha_c \in [0.4, 0.45]$) and selected the normalized profit of the solution such that the corresponding decision problem lied in the phase transition region (high $\text{TCC}_O$; $\alpha_p^* \in [0.6, 0.65]$) and in the over-constrained region (low $\text{TCC}_O$; $\alpha_p^* \in [0.85, 0.9]$). It is worth noting that the instance generation process did not produce instances in the under-constrained region ($\alpha_p^* \in [0.35 - 0.4]$). Again, we forced half of the instances selected in each of the bins in the phase transition region (high $\text{TCC}_O$) to be easier than the median, according to the *Gecode* propagations measure, and the other half to be harder than the median. We sampled a total of 18 instances, 12 with high $\text{TCC}_O$ and 6 with low $\text{TCC}_O$.

### S4 Appendix: Expected number of solution witnesses and the constrainedness of the solution space

In this section we characterize mathematically the expected *number of witnesses* (number of subsets of items that satisfy the constraints) of a random ensemble of instances of the knapsack decision problem. We start by presenting a formal definition of the knapsack problem and some alternative definitions that will then be useful to characterize the expected number of witnesses. Afterwards, we introduce the Dirichlet distribution, which will be used to model the distribution of random instances. We then characterize mathematically the expected number of witnesses. Finally, we show how the expected number of witnesses can be linked to the constrainedness of the solution space and how this is related to computational requirements.

**Defining the knapsack problem.** Let us recall the Knapsack Decision Problem:

**Definition 4** *Knapsack Decision Problem (KDP)*

*Let $n \in \mathbb{N}$, $c, p \in \mathbb{R}$ and $\psi = \{(w_1, v_1), ..., (w_n, v_n)\} \in \mathbb{R}^n \times \mathbb{R}^n$. A KDP is a mapping $K_n^{(c,p)} : \mathbb{R}^n \times \mathbb{R}^n \to \{0, 1\}$ such that:*

$$\bar{K}_n^{(c,p)}(\psi) = \begin{cases} 1 & \text{if } \exists A \subseteq \psi \text{ s.t. } \sum_{(w_i,v_i)\in A} w_i \leq c \text{ and } \sum_{(w_i,v_i)\in A} v_i \geq p \\ 0 & \text{otherwise} \end{cases}$$

We are interested in the KDP; however, to analyze the characteristics of the problem we investigate an analogous version of the problem that relates KDP to the normalized capacity ($\alpha_c = \frac{c}{\sum_{i=1}^{n} w_i}$) and normalized profit ($\alpha_p = \frac{p}{\sum_{i=1}^{n} v_i}$). We call this the normalized knapsack decision problem (NKDP), which is defined on a simplex:

**Definition 5** *m-simplex*

$$\triangle_m = \left\{(x_1, \ldots, x_{m+1}) \in \mathbb{R}^{m+1} | \sum_{i=1}^{m} x_i = 1 \text{ and } x_i \geq 0 \ \forall i \right\}$$

**Definition 6** *Normalized Knapsack Decision Problem (NKDP)*

*Let $n \in \mathbb{N}$ and $\alpha_c, \alpha_p \in H = \{x \in \mathbb{R} | 0 \leq x \leq 1\}$. A NKDP is a mapping $K_n^{(\alpha_c,\alpha_p)} : \triangle_{n-1} \times \triangle_{n-1} \to \{0, 1\}$ such that for any $\psi = \{(w_1, v_1), ..., (w_n, v_n)\} \in \triangle_{n-1} \times \triangle_{n-1}$:*

$$K_n^{(\alpha_c,\alpha_p)}(\psi) = \begin{cases} 1 & \text{if } \exists A \subseteq \psi \text{ s.t. } \sum_{(w_i,v_i)\in A} w_i \leq \alpha_c \text{ and } \sum_{(w_i,v_i)\in A} v_i \geq \alpha_p \\ 0 & \text{otherwise} \end{cases}$$

There is a correspondence between Definitions 4 and 6 by normalizing the weights and values of the former. Explicitly, if $\psi = \{(w_1, v_1), ..., (w_n, v_n)\} \in \mathbb{R}^n \times \mathbb{R}^n$ and $c, p \in \mathbb{R}$, we get that:

$$\bar{K}_n^{(c,p)}(\psi) = K_n^{(\alpha_c,\alpha_p)}(\hat{\psi})$$

where $\hat{\psi} = \{(\hat{w}_1, \hat{v}_1), ..., (\hat{w}_n, \hat{v}_n)\}$ with $\hat{w}_i = w_i / \sum_{i=1}^n w_i$, $\hat{v}_i = v_i / \sum_{i=1}^n v_i$, $\alpha_c = c / \sum_{i=1}^n w_i$ and $\alpha_p = p / \sum_{i=1}^n v_i$.

To simplify notation we now introduce an alternative version of Definition 6:

**Definition 7** *Normalized Knapsack Decision Problem (NKDP): Analogous Definition*

*Let $n \in \mathbb{N}$ and $\alpha_c, \alpha_p \in H = \{x \in \mathbb{R} | 0 \leq x \leq 1\}$. A NKDP is a mapping $K_n^{(\alpha_c,\alpha_p)} : \triangle_{n-1} \times \triangle_{n-1} \to \{0, 1\}$ such that for any $\psi = \{(w_1, v_1), ..., (w_n, v_n)\} \in \triangle_{n-1} \times \triangle_{n-1}$:*

$$K_n^{(\alpha_c,\alpha_p)}(\psi) = \begin{cases} 1 & \text{if } \exists S \subseteq \{1, 2, ..., n\} \text{ s.t. } \sum_{i \in S} w_i \leq \alpha_c \text{ and } \sum_{i \in S} v_i \geq \alpha_p \\ 0 & \text{otherwise} \end{cases}$$

Based on Definition 7, we define a *subset specific* NKDP. This is defined as a NKDP in which the question is whether a specific subset of items (e.g. items one and three) satisfy the normalized profit ($\alpha_p$) and normalized capacity ($\alpha_c$) constraints. ssNKDDP is defined for every set $S$ in the power set $\mathcal{P}(\{1, 2, ..., n\})$.

**Definition 8** *Subset Specific Normalized Knapsack Decision Problem (ssNKDP)*

*Let $n \in \mathbb{N}$, $\alpha_c, \alpha_p \in H = \{x \in \mathbb{R} | 0 \leq x \leq 1\}$ and $S \in \mathcal{P}(\{1, 2, ..., n\})$. A ssNKDP is a mapping $K_n^{(\alpha_c,\alpha_p,S)} : \triangle_{n-1} \times \triangle_{n-1} \to \{0, 1\}$ such that for any $\psi = \{(w_1, v_1), ..., (w_n, v_n)\} \in \triangle_{n-1} \times \triangle_{n-1}$:*

$$K_n^{(\alpha_c,\alpha_p,S)}(\psi) = \begin{cases} 1 & \text{if } \sum_{i \in S} w_i \leq \alpha_c \text{ and } \sum_{i \in S} v_i \geq \alpha_p \\ 0 & \text{otherwise} \end{cases}$$

**Sampling instances: The Dirichlet distribution.** We now turn our attention to the process of generation of random instances. We focus our attention on the Dirichlet probability distribution, which has been widely studied and is particularly suited to describe distributions over a simplex. This will allow us to characterize the expected number of witnesses of the NKDP in the following section. In this section we define the Dirichlet distribution and present some relevant properties.

**Definition 9** *Dirichlet Distribution*

*A random vector $(X_1, \ldots, X_m) \in \triangle_{m-1}$ is said to follow a Dirichlet distribution with parameters $\boldsymbol{\alpha} = (\alpha_1, \ldots, \alpha_m) \in (\mathbb{R}^+)^m$*

$$(X_1, \ldots, X_m) \sim Dir(\alpha_1, \ldots, \alpha_m)$$

*if the density of $(X_1, \ldots, X_{m-1})$ is*

$$\frac{\Gamma(\sum_{i=1}^m \alpha_i)}{\prod_{i=1}^m \Gamma(\alpha_i)} \prod_{i=1}^m X_i^{\alpha_i - 1}$$

**Lemma 1** *Additive Property*

*Let $m \in \mathbb{N}$ and $S \subseteq \{1, 2, \ldots, m\}$. If $(X_1, \ldots, X_m) \sim Dir(\alpha_1, \ldots, \alpha_m)$ then*

$$\left( \sum_{i \in S} X_i, \boldsymbol{X}_{-S} \right) \sim Dir\left( \sum_{i \in S} \alpha_i, \boldsymbol{\alpha}_{-S} \right)$$

**Lemma 2** *Marginal Distribution*

*If $(X_1, \ldots, X_m) \sim Dir(\alpha_1, \ldots, \alpha_m)$ then*

$$X_k \sim Beta\left( \alpha_k, \left( \sum_{i=1}^m \alpha_i \right) - \alpha_k \right)$$

**Corollary 1** *Let $m \in \mathbb{N}$ and $S \subseteq \{1, 2, \ldots, m\}$. If $(X_1, \ldots, X_m) \sim Dir(\alpha_1, \ldots, \alpha_m)$ then*

$$\sum_{i \in S} X_i \sim Beta\left( \sum_{i \in S} \alpha_i, \sum_{i \notin S} \alpha_i \right)$$

Let us turn back now to the knapsack problem and characterize the random generation process of instances. We sample weights and values independently from two Dirichlet distributions:

$$(w_1, \ldots, w_n) \sim Dir(\alpha, \ldots, \alpha)$$

$$(v_1, \ldots, v_n) \sim Dir(\beta, \ldots, \beta)$$

where $\alpha, \beta \in \mathbb{R}^+$. For simplicity we will restrict ourselves to the case where the $n$ weights and $n$ values are both sampled uniformly from the $(n-1)$-simplex:

$$(w_1, \ldots, w_n) \sim Dir(1, \ldots, 1)$$

$$(v_1, \ldots, v_n) \sim Dir(1, \ldots, 1)$$

The uniform case can be generalized easily to any other values of $\alpha$ and $\beta$.

**Expected number of witnesses.** Our aim is to characterize the expected number of witnesses of the NKDP when we introduce randomness in the selection of the items $\psi \in \triangle_{n-1} \times \triangle_{n-1}$. In order to do this we define first the random variable that captures the number of witnesses of a ssNKDP:

**Definition 10** *Number of Witnesses of a ssNKDP*
*Let* $X_n^{(\alpha_c,\alpha_p,S)} : \triangle_{n-1} \times \triangle_{n-1} \to \mathbb{N}$ *such that*

$$X_n^{(\alpha_c,\alpha_p,S)}(\psi) = K_n^{(\alpha_c,\alpha_p,S)}(\psi)$$

We now define the random variable of the number of witnesses of the NKDP as the sum of $X_n^{(\alpha_c,\alpha_p,S)}(\psi)$ over all possible subsets $S \in \mathcal{P}\{1,\dots,n\}$:

**Definition 11** *Number of Witnesses of a NKDP*
*Let* $X_n^{(\alpha_c,\alpha_p)} : \triangle_{n-1} \times \triangle_{n-1} \to \mathbb{N}$ *such that*

$$X_n^{(\alpha_c,\alpha_p)}(\psi) = \sum_{S\in\mathcal{P}(\{1,\dots,n\})} X_n^{(\alpha_c,\alpha_p,S)}(\psi)$$

In order to calculate the expected value of $X_n^{(\alpha_c,\alpha_p)}(\psi)$ it suffices to find the expected value of $X_n^{(\alpha_c,\alpha_p,S)}(\psi)$:

$$E\Big[X_n^{(\alpha_c,\alpha_p)}(\psi)\Big] = E\Bigg[\sum_{S\in\mathcal{P}(\{1,\dots,n\})} X_n^{(\alpha_c,\alpha_p,S)}(\psi)\Bigg] = \sum_{S\in\mathcal{P}(\{1,\dots,n\})} E\Big[X_n^{(\alpha_c,\alpha_p,S)}(\psi)\Big]$$

Given that $X_n^{(\alpha_c,\alpha_p,S)}(\psi) \in \{0,1\}$ we obtain that:

$$E\Big[X_n^{(\alpha_c,\alpha_p,S)}(\psi)\Big] = 1 \times P\Big(\sum_{i\in S} w_i \le \alpha_c \wedge \sum_{i\in S} v_i \ge \alpha_p\Big)$$

for an arbitrary $S \subseteq \{1,\dots,n\}$ and arbitrary $\alpha_c, \alpha_p \in H = \{x \in \mathbb{R} | 0 \le x \le 1\}$.

Given that the weights are sampled independently, we have

$$\begin{aligned} P\Big(\sum_{i\in S} w_i \le \alpha_c \wedge \sum_{i\in S} v_i \ge \alpha_p\Big) &= P\Big(\sum_{i\in S} w_i \le \alpha_c\Big) P\Big(\sum_{i\in S} v_i \ge \alpha_p\Big) \\ &= P\Big(\sum_{i\in S} w_i \le \alpha_c\Big)\Bigg(1 - P\Big(\sum_{i\in S} v_i \le \alpha_p\Big)\Bigg) \end{aligned}$$

Additionally, by Corollary 1, we can conclude that

$$\sum_{i\in S} w_i \sim Beta(|S|, n-|S|)$$

$$\sum_{i\in S} v_i \sim Beta(|S|, n-|S|)$$

where $|S|$ is the cardinality of the set. Therefore, if we denote the cumulative distribution of $Y \sim Beta(p,q)$ by $I_y(p,q)$, we get that

$$P\Big(\sum_{i\in S} w_i \le \alpha_c \wedge \sum_{i\in S} v_i \ge \alpha_p\Big) = I_{\alpha_c}\big(|S|, n-|S|\big)\big(1 - I_{\alpha_p}(|S|, n-|S|)\big).$$

Thus

$$\begin{aligned} E\Big[X_n^{(\alpha_c,\alpha_p)}(\psi)\Big] &= \sum_{S\in\mathcal{P}(\{1,\dots,n\})} I_{\alpha_c}\big(|S|, n-|S|\big)\big(1 - I_{\alpha_p}(|S|, n-|S|)\big) \\ &= \sum_{j=1}^{n} \binom{n}{j} I_{\alpha_c}\big(j, n-j\big)\big(1 - I_{\alpha_p}(j, n-j)\big). \end{aligned}$$

We summarize the above in the following result.

**Result 1** *NKDP expected number of witnesses*
*Let* $n \in \mathbb{N}$ *and* $\alpha_c, \alpha_p \in H = \{x \in \mathbb{R} | 0 \le x \le 1\}$. *If weights are sampled independently from values from the following distributions:* $(w_i, \dots, w_n) \sim Dir(1,\dots,1)$ *and* $(v_i,\dots,v_n) \sim Dir(1,\dots,1)$, *then the expected number of witnesses for the NKDP is given by*

$$E\Big[X_n^{(\alpha_c,\alpha_p)}(\psi)\Big] = \sum_{j=1}^{n} \binom{n}{j} I_{\alpha_c}\big(j, n-j\big)\big(1 - I_{\alpha_p}(j, n-j)\big)$$

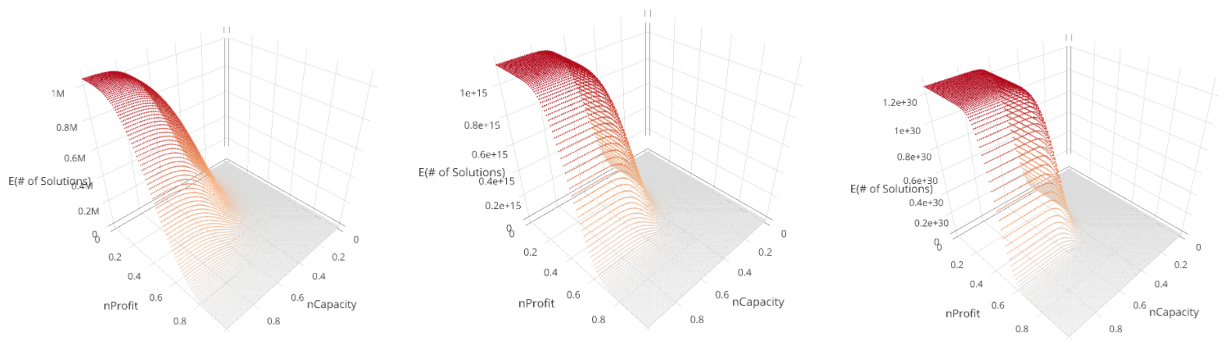


**(a)** n=20 **(b)** n=50 **(c)** n=100

**Fig. 1. Expected number of witnesses in the knapsack decision problem for different number of items (n).** Weights and values are sampled from independent uniform Dirichlet distributions. Equivalently, each value and weight is sampled from an independent $Gamma(1,1)$ distribution and then normalized with respect to the sum of values and weights, respectively.

In order to relate this result to the KDP we need to find a way of mapping a distribution of weights and values in $\mathbb{R}^n$ to a distribution in $\triangle_{n-1}$ and, in particular, to the Dirichlet distribution. Fortunately, every Dirichlet distribution can be constructed from independent Gamma distributions:

**Lemma 3** *$(X_1, \ldots, X_m) \sim Dir(\alpha_1, \ldots, \alpha_m)$ if and only if*

$$X_i \sim \frac{Y_i}{\sum_{i=1}^{m} Y_i} \text{ for } i = 1, \ldots, n$$

*where $Y_i \sim Gamma(\alpha_i, 1)$ and $\{Y_i\}_{i=1}^m$ are mutually independent.*

This implies the following result for KDP:

**Result 2** *KDP Expected number of witnesses*
*Let $n \in \mathbb{N}$ and $\alpha_c, \alpha_p \in H = \{x \in \mathbb{R} | 0 \le x \le 1\}$. If $w_i \sim Gamma(1,1)$ and $v_i \sim Gamma(1,1)$ are all mutually independent then*

$$E\left[\bar{X}_n^{(\alpha_c,\alpha_p)}(\psi)\right] = \sum_{j=1}^{n} \binom{n}{j} I_{\alpha_c}\big(j, n-j\big)\big(1 - I_{\alpha_p}(j, n-j)\big)$$

*where $\bar{X}_n^{(\alpha_c,\alpha_p)}(\psi)$ is defined for the KDP analogously to $X_n^{(\alpha_c,\alpha_p)}(\psi)$ for the NKDP (11).*

Using these results it is possible to calculate the expected number of witnesses for each $\alpha_c, \alpha_p \in H = \{x \in \mathbb{R} | 0 \le x \le 1\}$. We plot these values for different number of items (Fig 1).

**Constrainedness of the solution space, satisfiability phase transition and computational requirements.** The expected number of witnesses is tightly connected to the to the satisfiability phase transition. In particular, it has been suggested that the same parameters that characterizes the phase transition (i.e $\alpha_c$ and $\alpha_p$) characterize the expected number of witnesses. This has already been shown to apply to graph coloring, number partitioning and the boolean satisfiability problem (SAT) (5). In the previous section we showed further support for this claim by showing that, like the phase transition, the expected number of witnesses (for a fixed number of items $n$) is characterized by the same parameters, namely the normalized profit $\alpha_p$ and the normalized capacity $\alpha_c$. This suggests that the parameters related to the satisfiability phase transition in a computational problem can be found by deriving the analytical expression of the expected number of witnesses.

Notably, Gent et al. (5) propose that the parameter $\kappa$, which depends on the expected number of witnesses, characterizes the constrainedness of search. Explicitly,

$$\kappa = 1 - \frac{log_2(\text{Expected Number of witnesses})}{log_2(|states|)}$$

where *states* is the total state space of the problem.

We explore how the $\kappa$-parameter is related to the satisfiability phase transition and to computational requirements in the KDP. In order to do this, we first calculate $\kappa$. Note that in the knapsack, the states corresponds to all of the possible subsets of items (i.e. $S \in \mathcal{P}(\{1, \ldots, n\})$). In particular, we have that

$$|states| = |\mathcal{P}(\{1, \ldots, n\})| = 2^n$$

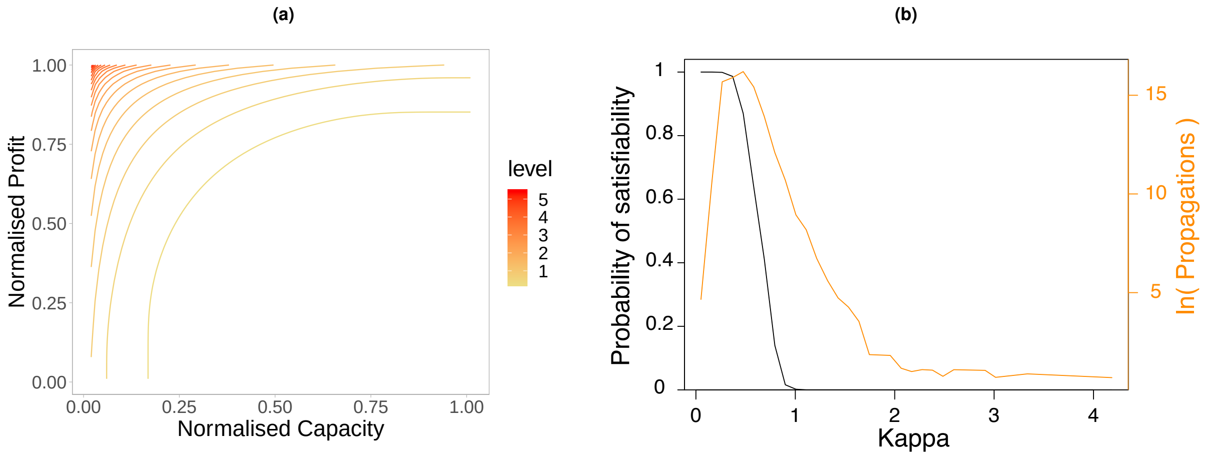


**Fig. 2. $\kappa$ for the Knapsack Decision Problem with 50 items.** Weights and values were independently sample from a $Gamma(1,1)$ distribution. **(a) $\kappa$ levels in the $\alpha_c \times \alpha_p$ space. (b) $\kappa$, satisfiability and computational requirements**. Satisfiability probability (probability of the existence of at least one witness; left axis) and the number propagations (right axis).

therefore, for the knapsack decision problem

$$\begin{aligned}\kappa &= 1 - \frac{log_2\left(E\left[\bar{X}_n^{(\alpha_c,\alpha_p)}(\psi)\right]\right)}{n} \\ &= 1 - \frac{1}{n} log_2\left[\sum_{j=1}^{n} \binom{n}{j} I_{\alpha_c}(j, n-j)\left(1 - I_{\alpha_p}(j, n-j)\right)\right]\end{aligned}$$

To test whether $\kappa$ is related to the constrainedness of the solution spaces we sampled 62,500 instances of the knapsack decision problem with $n = 50$ items. We solved the instances and calculated the satisfiability probability across the $\alpha_c \times \alpha_p$ space. Additionally, we calculated the average computational time required to solve instances. We found that, in line with Gent et al. (5), $\kappa$ characterizes a phase transition in the satisfiability probability (Fig 2b). Furthermore, we found that the average computational requirements peak around the same value of $\kappa$ where the phase transition occurs (Fig 2b). This give support to the claim that the parameters related to the expected number of witnesses and the satisfiability phase transition are closely related to the constrainedness of the solution space and to the expected computational requirements of solving an instance of the problem.

### S5 Appendix: CANTAB tasks

Four tests from the Cambridge Neuropsychological Test Automated Battery (CANTAB®) are used to measure certain aspects of cognitive function such as working memory and strategy use.

**Reaction Time (RTI)** Five yellow circles are displayed at the top of the screen, whilst the participant must press and hold down a touchscreen button at the bottom of the screen. When a spot appears inside one of the yellow circles the participant must respond as quickly as possible by letting go of the button and touching the circle where the yellow spot had appeared. This is repeated for 30 trials.

**Paired Associates Learning (PAL)** Boxes are displayed on the screen and open one by one in a randomized order to reveal patterns hidden inside. The patterns are then displayed in the middle of the screen, one at a time, and the subject must touch the box where the pattern was originally located.

**Spatial Working Memory (SWM)** The test begins with colored boxes being shown on the screen. The aim of this test is that, by touching the boxes and using a process of elimination, the subject should find one 'token' in each of the boxes and use them to fill up an empty column on the right hand side of the screen. The computer will never hide a token in the same colored box, so once a token is found in a box the participant should not return to that box to look for another token.

**Spatial Span Task (SSP)** White squares briefly change color in a variable sequence. The participant must remember the sequence and then touch the squares in that same order. The sequence length increases through the test. There are up to 3 attempts at each sequence length and the test terminates if all three are failed.

1. Gecode Team, Gecode: Generic Constraint Development Environment (2006).
2. N Een, N Sörensson, An extensible SAT-solver (2003).
3. N Eén, N Sörensson, Translating Pseudo-Boolean Constraints into SAT. *J. on Satisf. Boolean Model. Comput.* **2**, 1–25 (2006).

4. N Nethercote, et al., MiniZinc: Towards A Standard CP Modelling Language in *Proceedings of the 13th International Conference on Principles and Practice of Constraint Programming.* pp. 529–543 (2007).
5. IP Gent, E MacIntyre, P Prosser, T Walsh, The constrainedness of search. *AAAI/IAAI* (1996).

**S1 Table. Human performance in the knapsack decision task. Logistic regressions with random intercept effects for participants relating the Accuracy on an instance and Trial Number (1), Typical-case Complexity (TCC) (2), TCC and the Satisfiability (3), Gecode Propagations (4), Minisat Decisions (5), Over-constrained and Under-constrained regions (6), Over-constrained and Phase Transition regions (7), TCC, the Number of Witnesses and Satisfiability (8) and Instance Complexity (IC) (9).**

| | Dependent Variable: Human Performance | | | | | | | | |
|---|---|---|---|---|---|---|---|---|---|
| | (1) | (2) | (3) | (4) | (5) | (6) | (7) | (8) | (9) |
| Trial Number | 0.005<br>(0.004)<br>$p = 0.196$ | | | | | | | | |
| Typical-case Complexity (TCC) | | $-1.327^{***}$<br>(0.161)<br>$p < 0.001$ | $-1.285^{***}$<br>(0.240)<br>$p < 0.001$ | | | | $-1.208^{***}$<br>(0.202)<br>$p < 0.001$ | $-1.339^{***}$<br>(0.225)<br>$p < 0.001$ | |
| Number of Witnesses | | | | | | | | $0.139^{***}$<br>(0.041)<br>$p = 0.001$ | |
| TCC:Number of Witnesses | | | | | | | | $0.448^{***}$<br>(0.101)<br>$p < 0.001$ | |
| Satisfiability | | | $-0.250$<br>(0.271)<br>$p = 0.355$ | | | | | $-1.427^{***}$<br>(0.243)<br>$p < 0.001$ | |
| TCC:Satisfiability | | | $-0.084$<br>(0.323)<br>$p = 0.796$ | | | | | | |
| Gecode Propagations (scaled) | | | | $-0.362^{***}$<br>(0.066)<br>$p < 0.001$ | | | | | |
| Minisat Decisions | | | | | $-0.022$<br>(0.026)<br>$p = 0.395$ | | | | |
| Over-constrained | | | | | | $1.459^{***}$<br>(0.220)<br>$p < 0.001$ | 0.250<br>(0.271)<br>$p = 0.355$ | | |
| Under-constrained | | | | | | $1.208^{***}$<br>(0.202)<br>$p < 0.001$ | | | |
| $IC^{0.01}$ | | | | | | | | | $2.829^{***}$<br>(0.544)<br>$p < 0.001$ |
| Constant | $1.516^{***}$<br>(0.150) | $2.451^{***}$<br>(0.167) | $2.584^{***}$<br>(0.224) | $1.689^{***}$<br>(0.118) | $1.714^{***}$<br>(0.142) | $1.125^{***}$<br>(0.130) | $2.333^{***}$<br>(0.206) | $2.627^{***}$<br>(0.220) | $-1.117^{**}$<br>(0.544) |
| Observations | 1,427 | 1,427 | 1,427 | 1,427 | 1,427 | 1,427 | 1,427 | 1,427 | 1,427 |
| Log Likelihood | -639.577 | -602.372 | -600.149 | -626.120 | -640.066 | -601.946 | -601.946 | -581.201 | -624.276 |
| Akaike Inf. Crit. | 1,285.153 | 1,210.744 | 1,210.299 | 1,258.240 | 1,286.131 | 1,211.891 | 1,211.891 | 1,174.402 | 1,254.552 |
| Bayesian Inf. Crit. | 1,300.943 | 1,226.534 | 1,236.615 | 1,274.030 | 1,301.921 | 1,232.944 | 1,232.944 | 1,205.982 | 1,270.342 |

*Note:* $^{*}$p<0.1; $^{**}$p<0.05; $^{***}$p<0.01

**S2 Table. Computational performance in the knapsack optimization task. Logistic regressions with random intercept effects for participants relating Computational Performance on an instance and Trial Number (1), Optimisation Typical-case Complexity ($TCC_O$) (2), $TCC_O$ and the Time Spent (3), Gecode Propagations (4), Minisat Decisions (5), Sahni-K (6), and Time Spent (7).**

| | Dependent Variable | | | | | | |
|---|---|---|---|---|---|---|---|
| | Computational performance | | | | | | |
| | (1) | (2) | (3) | (4) | (5) | (6) | (7) |
| Trial Number | $0.012$<br>$(0.030)$<br>$p = 0.683$ | | | | | | |
| Time Spent (scaled) | | | $-0.089$<br>$(0.735)$<br>$p = 0.905$ | | | | |
| Typical-case Complexity ($TCC_O$) | | $-2.175^{***}$<br>$(0.531)$<br>$p < 0.001$ | $-2.420^{***}$<br>$(0.817)$<br>$p = 0.004$ | | | | |
| Time Spent (scaled): $TCC_O$ | | | $-0.709$<br>$(0.760)$<br>$p = 0.352$ | | | | |
| Gecode Propagations | | | | $-0.108^{***}$<br>$(0.022)$<br>$p < 0.001$ | | | |
| Minisat Decisions | | | | | $-0.026$<br>$(0.018)$<br>$p = 0.157$ | | |
| Sahni-K | | | | | | $-1.333^{***}$<br>$(0.193)$<br>$p < 0.001$ | |
| Time Spent | | | | | | | $-0.072^{***}$<br>$(0.017)$<br>$p < 0.001$ |
| Constant | $1.512^{***}$<br>$(0.261)$ | $3.359^{***}$<br>$(0.509)$ | $3.914^{***}$<br>$(0.796)$ | $4.230^{***}$<br>$(0.593)$ | $2.124^{***}$<br>$(0.404)$ | $2.310^{***}$<br>$(0.219)$ | $4.975^{***}$<br>$(0.865)$ |
| Observations | 358 | 358 | 305 | 358 | 358 | 358 | 305 |
| Log Likelihood | -161.752 | -147.622 | -115.185 | -149.815 | -160.808 | -135.816 | -124.754 |
| Akaike Inf. Crit. | 329.504 | 301.245 | 240.369 | 305.630 | 327.615 | 277.633 | 255.509 |
| Bayesian Inf. Crit. | 341.146 | 312.887 | 258.971 | 317.272 | 339.257 | 289.274 | 266.670 |

*Note:* $^{*}$p<0.1; $^{**}$p<0.05; $^{***}$p<0.01

**S3 Table. Effort in the knapsack optimization task. Linear regressions with random intercept effects for participants relating Time Spent on an instance and Trial Number (1), Optimization Typical-case Complexity ($TCC_O$) (2), Gecode Propagations (3), Minisat Decisions (4), Sahni-K (5), $TCC_O$ together with computational performance (6), and $TCC_O$ together with Sahni-K.**

| | Dependent Variable | | | | | | |
|---|---|---|---|---|---|---|---|
| | Time Spent | | | | | | |
| | (1) | (2) | (3) | (4) | (5) | (6) | (7) |
| Trial Number | 0.095<br>(0.135)<br>$p = 0.483$ | | | | | | |
| Typical-case Complexity ($TCC_O$) | | $10.114^{***}$<br>(1.236)<br>$p < 0.001$ | | | | $15.161^{**}$<br>(7.261)<br>$p = 0.037$ | $9.535^{***}$<br>(1.366)<br>$p < 0.001$ |
| Gecode Propagations | | | $0.438^{***}$<br>(0.100)<br>$p < 0.001$ | | | | |
| Minisat Decisions | | | | $0.276^{***}$<br>(0.080)<br>$p = 0.001$ | | | |
| Sahni-K | | | | | $3.125^{***}$<br>(0.950)<br>$p = 0.001$ | | 1.489<br>(2.687)<br>$p = 0.580$ |
| Sahni-K:$TOC_O$ | | | | | | | 0.502<br>(2.844)<br>$p = 0.861$ |
| Computational Performance | | | | | | $-0.489$<br>(7.210)<br>$p = 0.946$ | |
| Computational Performance : $TCC_O$ | | | | | | $-6.784$<br>(7.374)<br>$p = 0.358$ | |
| Constant | $41.802^{***}$<br>(2.210) | $35.749^{***}$<br>(2.132) | $32.505^{***}$<br>(3.013) | $37.165^{***}$<br>(2.503) | $41.469^{***}$<br>(1.990) | $36.227^{***}$<br>(7.379) | $35.498^{***}$<br>(2.178) |
| Observations | 305 | 305 | 305 | 305 | 305 | 305 | 305 |
| Log Likelihood | -1,188.894 | -1,156.717 | -1,180.119 | -1,183.824 | -1,181.856 | -1,142.942 | -1,151.532 |
| Akaike Inf. Crit. | 2,385.788 | 2,321.433 | 2,368.238 | 2,375.647 | 2,371.712 | 2,297.885 | 2,315.065 |
| Bayesian Inf. Crit. | 2,400.670 | 2,336.314 | 2,383.119 | 2,390.529 | 2,386.593 | 2,320.207 | 2,337.386 |

*Note:* $^{*}$p<0.1; $^{**}$p<0.05; $^{***}$p<0.01

**S4 Table. Other measures of performance in the knapsack optimization task. Linear regressions with random intercept effects for participants relating Optimization Typical-case Complexity ($TCC_O$) to Economic Performance (1), and Item Performance (2).**

| | Dependent Variable | |
|---|---|---|
| | Economic Performance | Item Performance |
| | (1) | (2) |
| Typical-case Complexity ($TCC_O$) | $-0.015^{***}$ | $0.410^{***}$ |
| | $(0.004)$ | $(0.092)$ |
| | $p < 0.001$ | $p < 0.001$ |
| Constant | $0.999^{***}$ | $0.076$ |
| | $(0.003)$ | $(0.076)$ |
| Observations | 347 | 358 |
| Log Likelihood | 685.782 | -441.574 |
| Akaike Inf. Crit. | -1,363.564 | 891.147 |
| Bayesian Inf. Crit. | -1,348.166 | 906.669 |
| *Note:* | | $^{*}$p<0.1; $^{**}$p<0.05; $^{***}$p<0.01 |

**S5 Table. Time spent after each item selection in the knapsack optimization task. Linear regressions with random intercept effects for participants relating the Time Spent at each item selection with IC and TCC (2), and The number of item-subsets that perform better than the current selection of items. Each selection of items is associated with the number of item-subsets that satisfy the capacity constraint and have higher sum of values than the current selection (1).**

| | Dependent Variable | |
|---|---|---|
| | Time spent at each item selection | |
| | (1) | (2) |
| Number of item-subsets that perform better | $-0.460^{***}$ | |
| | $(0.022)$ | |
| | $p < 0.001$ | |
| IC | | $-17.276^{***}$ |
| | | $(0.807)$ |
| | | $p < 0.001$ |
| TCC | | $-0.564$ |
| | | $(0.445)$ |
| | | $p = 0.206$ |
| Constant | $13.240^{***}$ | $14.053^{***}$ |
| | $(0.531)$ | $(0.629)$ |
| | $p < 0.001$ | $p < 0.001$ |
| Observations | 1781 | 1,781 |
| Log Likelihood | -6,410.726 | -6,392.535 |
| Akaike Inf. Crit. | 12,829.450 | 12,795.070 |
| Bayesian Inf. Crit. | 12,851.390 | 12,822.500 |

*Note:* $^{*}$p<0.1; $^{**}$p<0.05; $^{***}$p<0.01

**S6 Table. Pearson correlation between performance in the knapsack task and cognitive abilities. Performance in the knapsack decision task is characterized by Accuracy and in the knapsack optimization task is characterized by Computational Performance. The cognitive abilities measured used were Episodic Memory (PALFAMS28), Working Memory (SSPFSL), Strategy Use (SWMS) and Spatial Working Memory (weighted SWMTE, with errors on easier tasks being weighted more). Results are shown without multiple comparisons correction.**

| Task | Knapsack Decision | Knapsack Optimization |
|---|---|---|
| Mental Arithmetic | $0.022$ | $0.359$ |
| | $(0.236)$ | $(0.220)$ |
| | $p = 0.926$ | $p = 0.120$ |
| Episodic Memory | $-0.113$ | $0.191$ |
| | $(0.234)$ | $(0.231)$ |
| | $p = 0.637$ | $p = 0.420$ |
| Working Memory | $-0.019$ | $0.210$ |
| | $(0.236)$ | $(0.230)$ |
| | $p = 0.936$ | $p = 0.374$ |
| Strategy Use | $-0.342$ | $-0.351$ |
| | $(0.221)$ | $(0.221)$ |
| | $p = 0.140$ | $p = 0.129$ |
| Spatial Working Memory | $-0.165$ | $-0.325$ |
| | $(0.232)$ | $(0.222)$ |
| | $p = 0.487$ | $p = 0.162$ |
| Degrees of Freedom | 18 | 18 |

**S7 Table. Human accuracy and Complexity in the Knapsack Decision Task.** $R^2$ **and AIC model fit values of alternative models. Each of the models predicts average accuracy of an instance based on the complexity of the instance. 2 instances were identified as outliers and excluded from the analysis. The total number of observations in each model is** $n = 70$**.**

| Model | $R^2$ | AIC |
|---|---|---|
| $accuracy = \beta_0 + \beta_{IC} \times IC$ | 0.296 | −84.786 |
| $accuracy = \beta_0 + \beta_{IC} \times IC^{0.5}$ | 0.433 | −99.961 |
| $accuracy = \beta_0 + \beta_{IC} \times IC^{0.1}$ | 0.529 | −112.956 |
| $accuracy = \beta_0 + \beta_{IC} \times IC^{0.05}$ | 0.537 | −114.083 |
| $accuracy = \beta_0 + \beta_{IC} \times IC^{0.01}$ | 0.542 | −114.834 |
| $accuracy = \beta_0 + \beta_{IC} \times ln(IC)$ | 0.328 | −88.018 |
| $accuracy = \beta_0 + \beta_{IC} \times log(IC)$ | 0.328 | −88.018 |
| $accuracy = \beta_0 + \beta_{TCC} \times TCC$ | 0.213 | −77.053 |
